# Reliable population signal of subjective economic value from unreliable neurons in primate orbitofrontal cortex

**DOI:** 10.1101/2021.11.13.468353

**Authors:** Simone Ferrari-Toniolo, Wolfram Schultz

**Affiliations:** Department of Physiology, Development and Neuroscience, University of Cambridge, Cambridge, UK

**Keywords:** Reward, choice, utility, probability weighting, reliability

## Abstract

Behavior-related neuronal signals often vary between neurons. Despite the unreliability of individual neurons, brains are able to accurately represent and drive behavior. The notion may also apply to economic (‘value-based’) choices and the underlying reward signals. Reward value is subjective and can be defined by nonlinear weighting of magnitude (utility) and probability. Using a wide variety of reward magnitude and probability, we assessed subjective reward value at choice indifference between safe and risky rewards as prescribed by the continuity axiom that provides stringent criteria for meaningful choice. We found that individual neurons in the orbitofrontal cortex (OFC) of monkeys carry unreliable and heterogeneous neuronal signals for subjective value that largely fails to match the animal’s choice. However, the averaged neuronal signals matched well the animals’ choices, suggesting reliable subjective economic value encoding by the observed population of unreliable neurons.

**Highlights:**

- Different from widely held views, reliable neuronal information processing may not require reliable processors.
- Neurons in monkey orbitofrontal cortex (OFC) process reward magnitude and probability heterogeneously and unreliably.
- Despite unreliable neuronal processing, OFC population activity codes choices reliably.
- Reliability systems performance from unreliable elements seems to be a broad feature of neuronal reward coding.

**In brief:** Using stringent concepts of behavioral choice, Ferrari-Toniolo and Schultz describe unreliable individual reward neurons in monkey orbitofrontal cortex whose activity combines to a reliable population code for economic choice.

## INTRODUCTION

The construction of reliable systems from unreliable components has been recognized as a problem as soon as computing systems became formalized. The ‘Heuristic Argument’ states that the same information is simultaneously fed into several noisy processing elements and the accuracy results from averaging the noisy inputs (von Neumann, 1956). The brain, with its neurons as processing elements, may constitute a biological instantiation of such an automaton. The diverse anatomical and physiological nature of neurons allows heterogeneous representations of incoming information that can be summed in specific neuronal populations to obtain correct behaviors. For example, while reward prediction error responses of dopamine neurons are heterogeneous across individual neurons (Dabney et al., 2020), stimulation of the populations of dopamine neurons elicits reliable behavioral and neuronal learning (Stauffer et al., 2016; Saunders et al., 2018). Thus, a summing of unreliable neuronal responses may result in reliable, behavior-matching performance of a neuronal population.

Economic value refers to the value of a reward that is specific for the individual decision maker. Thus, economic value is subjective and determines individual welfare and evolutionary fitness. Situation-specific assessments of subjective economic value can employ satiety and temporal discounting (Padoa-Schioppa and Assad, 2006; Kobayashi and Schultz, 2008), but a general definition of subjective value requires mathematical functions that are able to predict choices. Expected Utility Theory (EUT) provides such functions by describing how choices maximize utility: decision makers assign a numerical utility to each option and choose the option with the highest utility (von Neumann and Morgenstern, 1944). However, EUT fails to explain risky choices in some situations (Allais, 1953). Built on elements of EUT, Prospect Theory (PT) provides a more complete description of risky choice by defining subjective economic value as the combination of subjectively weighted reward magnitude (utility) and subjectively weighted reward probability (Kahneman and Tversky, 1979). With these properties, PT is currently the simplest theory for defining a general foundational variable for economic choice suitable for investigation in reward neurons (Camerer, 2008; O’Doherty and Camerer, 2015). We will use this specific, PT-derived definition of subjective economic value rather than the less complete definition as nonlinear utility (Stauffer et al., 2014) or from ad-hoc, situation-specific assessments of general subjective value during choices and temporal discounting (Padoa-Schioppa and Assad, 2006; Kobayashi and Schultz, 2008; Lak et al., 2014).

Neurons sensitive to reward magnitude and probability have been identified in the dopaminergic midbrain and in the orbitofrontal cortex (OFC) (Tremblay and Schultz, 1999; Fiorillo et al., 2003; Padoa-Schioppa and Assad, 2006; Padoa-Schioppa and Cai, 2011; Lak et al., 2014; Stauffer et al., 2014; Imalzumi et al., 2022). Dopamine neurons show nonlinear responses to reward magnitude, compatible with encoding subjectively weighted reward magnitude or, in other words, mathematically defined economic utility (Stauffer et al., 2014). However, a closer look at the dopamine signal revealed considerable heterogeneity among individual neurons and demonstrated a more reliable neuronal teaching signal when considering the whole distribution of the neuronal response (Dabney et al., 2022). When considering decision-making, OFC neurons code subjective value related to choice preferences, spontaneous satiation and temporal discounting (Tremblay and Schultz, 1999; Padoa-Schioppa and Assad, 2006; Kobayashi and Schultz, 2008). More recent examinations demonstrate OFC coding of a wide variety of decision variables (Hirokawa et al., 2019), including utility and weighted probability as proposed by PT (Imalzumi et al., 2022). It would now be interesting to know whether these various and heterogeneous OFC responses might reliably correspond to the choices of the animal.

The current study aimed to apply the question of reliable neuronal coding to the processing of reward value in one of the main reward structures of the brain, the OFC. We benefitted from knowledge acquired in previous reward studies that had described basic properties of reward and risk signals in monkey OFC neurons. The crucial variables determining reward value are subjectively weighted reward magnitude (utility) and subjectively weighted reward probability. We had previously determined the validity of these variables in behavioral tests of the stringent continuity axiom (Ferrari-Toniolo et al., 2021); axiom compliance demonstrates understanding of the stimuli and meaningful choice behavior (von Neumann and Morgenstern, 1944), and thus the meaningful processing of subjective value inferred from observable choice. Our study had also shown that PT, based on utility and weighted probability, provided better fits to the choices compared to the mean-variance approach (which is based on expected value and variance risk). On the basis of meaningful reward valuations demonstrated by the axiomatic behavioral tests, we found that neuronal value signals in individual OFC neurons only rarely reflected the animal’s choices correctly. By contrast, the averaged population signals from these unreliable OFC signals matched well the animals’ choices, including their risk attitude. These results advance our basic knowledge of neuronal decision mechanisms by demonstrating how unreliable value signals of individual OFC neurons converge into a reliable neuronal population code for subjective economic value.

## RESULTS

### Design and behavior

This study aimed to advance the existing knowledge of reward and risk signals in OFC neurons (Tremblay and Schultz, 1999; Padoa-Schioppa and Assad, 2006; Kobayashi et al., 2010; Hirokawa et al., 2019; Imalzumi et al., 2022) by investigating the reliability of neuronal processing of subjective economic value. Based on the continuity axiom of EUT whose compliance demonstrates meaningful choices (von Neumann and Morgenstern, 1944), we tested a wide variety of reward magnitudes and probabilities in a structured manner. Our preceding behavioral study had demonstrated compliance of monkeys’ choices with this axiom (Ferrari-Toniolo et al., 2021). These neuronal and behavioral foundations provided the basis for testing the reliability of neuronal processing of the two fundamental components of subjective reward value, utility and weighted probability that underly economic choice.

We used three rewards labelled A, B and C according to decreasing value. Each reward consisted of the same liquid but had a specific magnitude (*m*; measured in ml: reward A: 0.20–0.50 ml, reward B: 0.15–0.45 ml, reward C: 0 ml). Each reward was delivered with probability *p* = 1.0 (for consistency with standard economic nomenclature, these rewards would be labelled as ‘degenerate’ gambles with only one ‘safe’ outcome).

In each trial, the monkey chose between two options presented at pseudorandomly alternating fixed left and right positions (Figure 1A, B). One option was the safe reward B with a preset fixed magnitude. The alternative option was a probabilistic combination of the two safe rewards A and C, weighted by probabilities *p*_A_ and (1 – *p*_A_). The resulting gamble AC was defined as

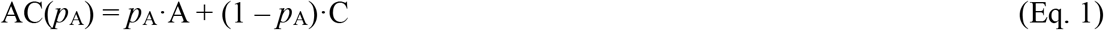

Thus, the AC combination was simply a gamble with magnitude *m* (equal to the magnitude of reward A) and probability *p* = *p*_A_ (in conventional notation (*m*_A_, *p*_A_)). We varied probability *p*_A_ to estimate choice functions for options AC vs. B (Figure 1C).

**Figure 1.**
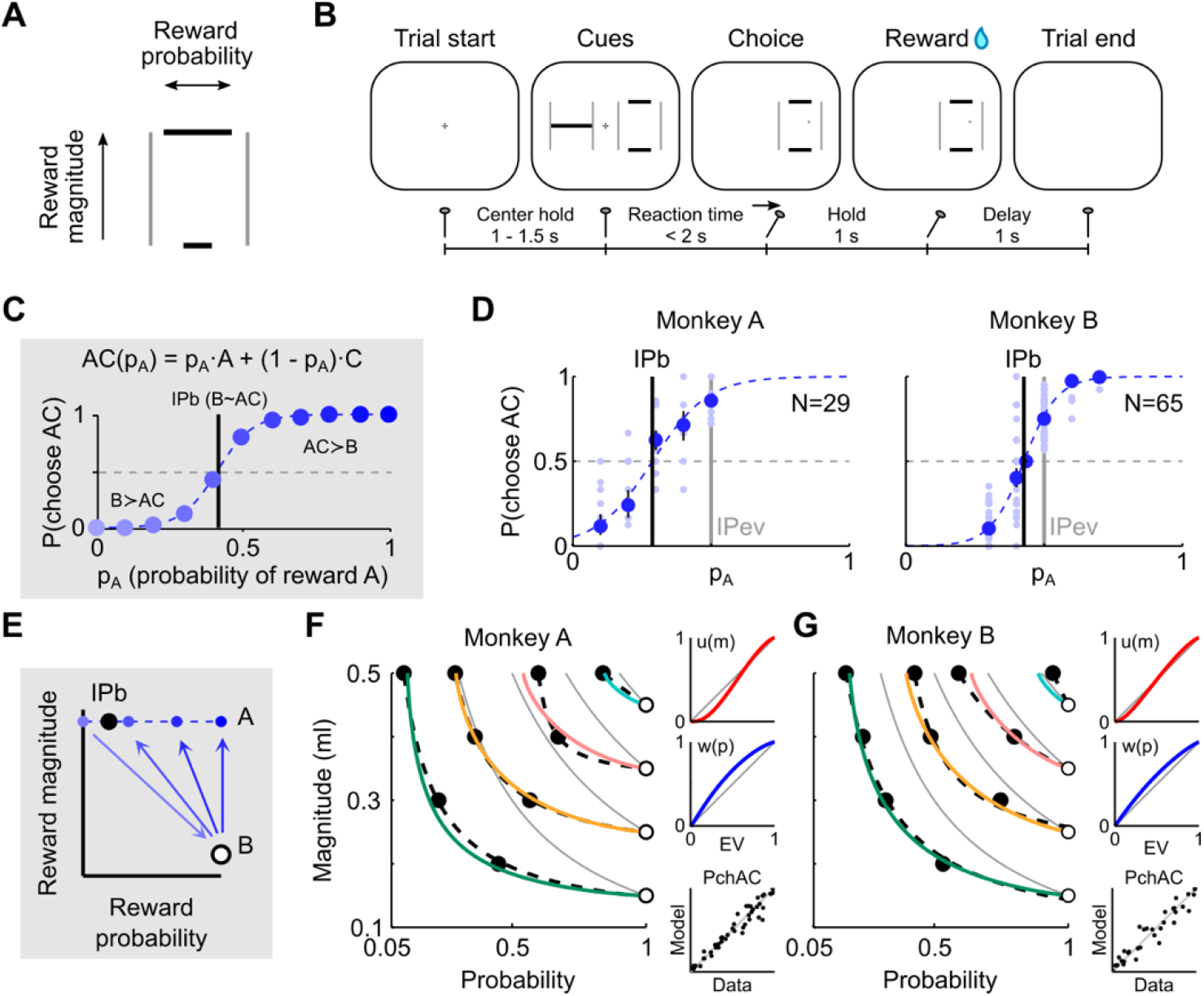
Utility and probability weighting functions describe risky choices. (A) Visual Cue. Each horizontal bar represents the magnitude (vertical position) and probability (horizontal width) of a possible outcome. Rewards varied between 0 and 0.5 ml. (B) Trial sequence. A monkey was presented with two choice options (Cues) and revealed its preference by moving a joystick toward one side. After 1 sec the reward was delivered, conditional on the reward distribution of the selected Cue. For neuronal recordings, only one of the two Cues was presented to the animal (zero-alternative task). (C) Scheme of subjective economic value estimation, based on the continuity axiom of EUT. In choices between a gamble AC (magnitude A with probability *p*_A_ and magnitude C with probability 1-*p*_A_) and safe option B, we estimated a behavioral indifference point (IPb). The IPb was defined as the probability (*p*_A_) for which the two options were equally frequently chosen and thus corresponded to a numerical value estimate for option B in relation to the combinatorial gamble AC. (D) Experimental results for one set of magnitudes {A = 0.50 ml, B = 0.25 ml, C = 0 ml}. Dots: probability of choosing option AC (pAC) in single sessions (light blue) and averages across sessions (dark blue) with 95% binomial confidence intervals (vertical bars). The softmax function *S(p_A_)* (dashed blue line) was fitted to the pAC to estimate the behavioral indifference point (IPb) as the *p*_A_ for which *S(p_A_)* = 0.5. IPev: *p*_A_ for which both options have same Expected Value (EV): *p*_A_ for which EV(B) = EV(AC). (E) Scheme of extended subjective economic value measurements. The procedure described in (C), represented in a two-dimensional magnitude-probability space. Arrows link the two presented options, pointing toward the preferred one. (F), (G) Indifference curves (ICs) and economic functions. Left: ICs in the magnitude-probability space were elicited by first measuring IPs for several combinations of gambles {A,B,C} (n = 9 in Monkey A, n = 10 in Monkey B); for a fixed B (open circle), all IPb’s (filled circles) had the same subjective economic value, thus belonging to the same IC (dashed line; hyperbolic functions (*m*(*p*) = c_1_ + c_2_ / (*p* – c_3_)) best-fitting the data points). The difference between ICs and EV-curves (gray lines: points with constant EV) quantifies the subjective valuation of choice options, indicating a risk seeking attitude (ICs shifted to the left) or a risk averse attitude (ICs shifted to the right). Model-based ICs (colored lines, Eq. 8) were estimated from all choices, for each monkey. Right: model’s utility function (*u*, top) and probability weighting function (*w*, middle) recovered from choices across all tests and sessions. The economic model robustly reconstructed preferences across all tests (bottom; PchAC: probability of choosing option AC; axes’ range: [0, 1]) (Monkey A: Pearson’s r = 0.97, P = 2.3e-30; Monkey B: r = 0.96, P = 2.8e-20). See also Figures S1 and S10.

We used observable behavioral choice to infer subjective economic value. For thoroughly testing the reliability of neuronal signals across the largest feasible variation of reward magnitude and probability, we relied on the continuity axiom as a conceptually solid and well-tested means for demonstrating meaningful choice. Using the axiom, we estimated subjective economic value from choice indifference as the reward probability *p*_A_ at which the variable composite gamble AC and the fixed gamble B were chosen equally frequently (choice indifference; P(choose each option) = 0.5) (Eq. 2). Tests at choice indifference are immune to slope differences of choice functions that might confound value estimation. We quantified preference (i.e., the probability of choosing the AC option) for different AC gambles, and fitted a softmax function to estimate the probability *p*_A_ at the behavioral choice indifference point (IPb) (Figure 1C) (Eq. 3). The estimated IPb represented the numeric subjective economic value of gamble B in relation to gambles A and C.

The animals made consistent choices that allowed us to estimate IPb’s in a single option-set {A,B,C} (Figure 1D), as well as in a variety of option-sets with varying gambles A and B. Importantly, at the behavioral choice indifference point IPb, the animal chose in half the trials the AC gamble with the physical lower mean reward and forwent the physical larger safe reward B, thus suggesting maximization of subjective economic value rather than physical value. We represented the results of these multiple-option-sets measurements in a two-dimensional magnitude-probability space, where each point corresponded to a gamble (Figure 1E). The full set of IPb’s identified a series of indifference curves (ICs) that connected gambles with equal subjective economic value in the magnitude-probability space (Figure 1F, G; dashed lines: hyperbolic curves fitted to the IPb’s). The distance between the ICs and the expected value (EV) curves (i.e., connected points with the same objective EV) highlighted the subjective nature of economic value, as revealed by choices. We fitted an economic model in which each gamble’s value was defined by a subjective utility function of reward magnitude *u(m)* and a subjective probability weighting function *w(p),* describing the nonlinear subjective weighting of reward magnitude (*m*) and probability (*p*), respectively (Eq. 4). Through a single set of fitted parameters, the economic model captured the behavioral preferences (Figure S1) and elicited ICs (Eq. 8) matching the measured IPb’s (Figure 1F, G; continuous lines).

These results demonstrate that the choice behavior in our task can be understood by assuming a combination of utility and probability weighting functions, in line with economic theory (see Ferrari-Toniolo et al., 2021 for more details).

### Neuronal magnitude and probability coding

We recorded extracellular activity from 838 single OFC neurons (Monkey A: 206 neurons, Monkey B: 632 neurons) in a zero-alternative version of the task in which the animal was presented with a quantitative stimulus predicting either the safe reward B or the gamble AC at a pseudorandomly alternating fixed left or right position, selected via a hand-held joystick-operated cursor. Both monkeys had a side bias, which we compensated by averaging the neuronal responses between left and right stimulus positions. Of the 838 neurons recorded in zero-alternative trials, 622 neurons (74%) showed significant responses to the presented Cue within the 400 ms analysis time-window (Figure 1B) (Monkey A: 149 neurons, Monkey B: 473 neurons) (P < 0.05; paired Wilcoxon test vs. pre-Cue baseline activity).

In an initial analysis that provided the baseline for this study, we identified single-neuron responses according to their reward magnitude and probability coding: neurons were only considered when their Cue responses increased or decreased significantly monotonically with both reward magnitude and probability in both safe reward B and gamble AC trials (P < 0.05; slopes bM · bP > 0, multiple linear regression, Eq. 9, indicating either increases with both variables (positive *m, p* coding) or decreases with both variables (inverse *m, p* coding). Of the 622 neurons with Cue responses, 166 neurons showed such response characteristics (Figure 2A-C) (Monkey A: 28 neurons, Monkey B: 138 neurons). The anatomical locations of all recorded neurons are shown in Figure S2. While these magnitude and probability coding neuronal responses served as stringent baseline, we describe below a separate analysis that compared more directly the neuronal with the behavioral subjective value measures, without imposing linear magnitude and probability coding (Eq. 12; see below).

**Figure 2.**
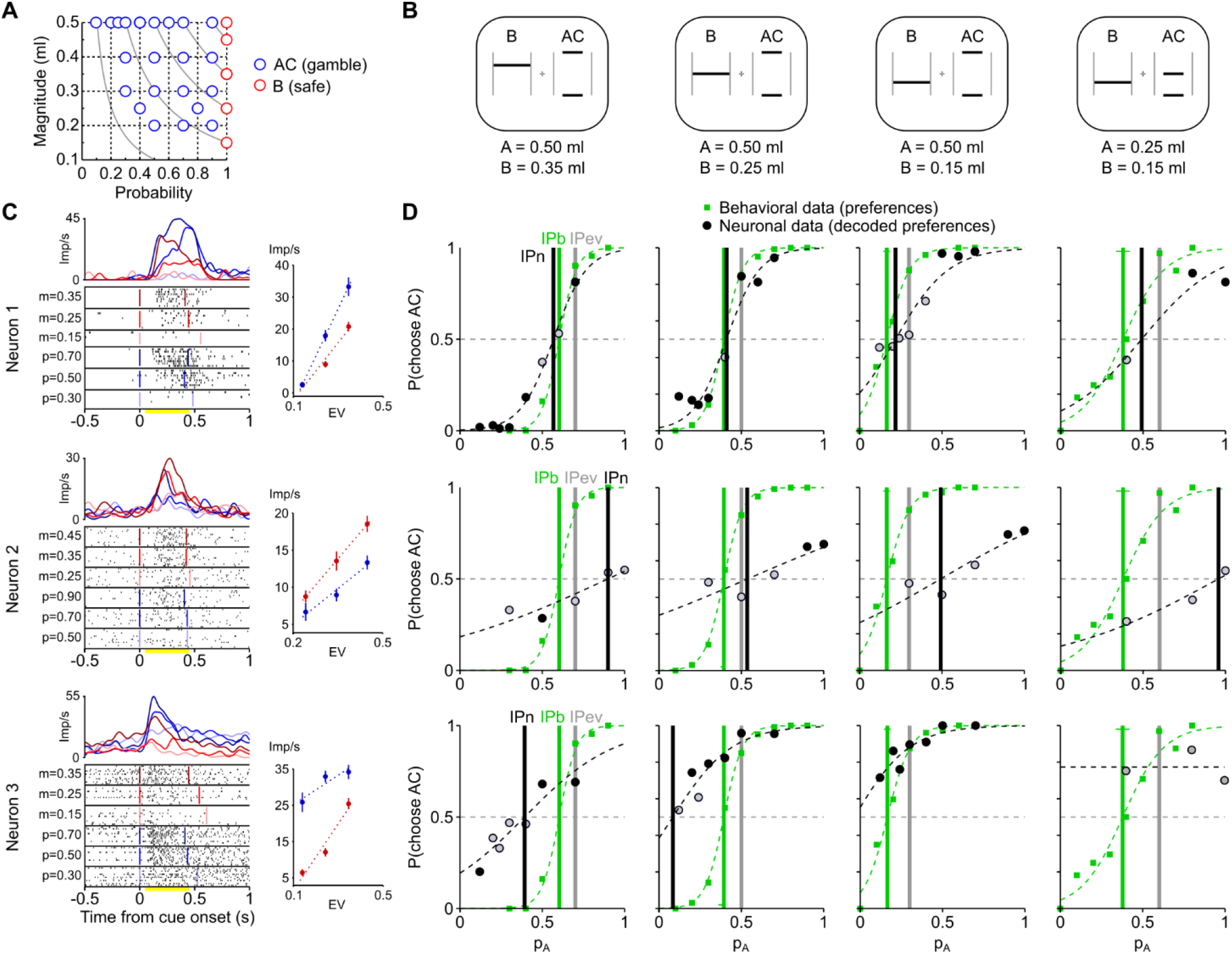
Heterogeneous subjective economic value coding in single neurons. (A) Set of gambles (circles) tested in neurons in a zero-alternative task, presenting them in pseudo-random alternation. For each neuron, the experimental test included 3 to 5 safe magnitudes (B) and 9 to 15 AC gambles (spanning at least 2 magnitude levels and 3 p_A_ probabilities). Gray lines: EV-curves. (B) Subjective economic value tests. Gamble Cues for four sample value measures. Each test is identified by the indicated combination of A and B magnitudes (with fixed C = 0 ml). For simplicity, only one AC Cue (p_A_ = 0.5) is shown. During neuronal recordings, only one option was presented to the monkey. (C) Raster plots and average responses for three example neurons recorded from Monkey B. Left: response to the option indicated on the y-axis (*m:* B magnitude, *p:* probability of A). Vertical lines indicate, for each trial class, the times of Cue presentation and the average behavioral reaction time. Top: spike density functions (gaussian-smoothed activity rates, kernel 30 ms) for the safe (red) or gamble (blue) option (darker colors for higher EV). Three safe options and three fixed-magnitude gambles (A = 0.5 ml) are represented here. Right: average (± SEM) activity rates computed in the analysis window (yellow bar in the raster plot). Linear fits (dotted lines) show sensitivity of neurons to both reward magnitude and probability. Imp/s: impulses per second. For neuronal responses in choice trials see Figure S3A. (D) Neuronal measure of subjective economic value. A neuronal indifference point (IPn, black vertical line) and the corresponding behavioral one (IPb, green vertical line) were computed for each test (i.e., options set). Each dot represents the probability of choosing the AC option (pAC) for a specific p_A_. The neuronal pACs (black) were decoded from the overlap in neuronal response distributions to the two choice options (see Methods and Figure S4); black-filled pACs: activity significantly different between the AC and B options (two-sample t-test, P < 0.05). Behavioral pAC (green): proportion of trials for which the AC option was chosen. IPev: p_A_ for which EV(B) = EV(AC). The IPn varied across neurons and the difference between IPn and IPb varied across tests, indicating that single neurons were not homogeneously encoding the options’ subjective economic value. For choice trial data see Figure S3B.

For comparison, we recorded from 319 single OFC neurons in choice trials (Monkey A: 191 neurons, Monkey B: 128 neurons), and 229 of these neurons (72%) responded significantly to the presented Cue within the 400 ms analysis time-window (Monkey A: 133 neurons, Monkey B: 96 neurons). Of these neurons, 89 neurons showed significantly monotonically increasing or decreasing responses with both reward magnitude and probability of the chosen option (Figure S3A) (39% of the 229 cue-responsive neurons recorded during choice; Monkey A: 54 neurons, Monkey B: 35 neurons; Eq. 9). With more formal testing, we found 64 neurons coding chosen value (28% of the 229 cue-responsive neurons; Monkey A: 36 neurons; Monkey B: 28; Eq. 10a-c). These neurons contrasted with 34 neurons that coded the value of only one of the choice objects (gamble AC or safe reward B) (Eq. 10d-e), an assessment that was possible due to their visual distinction (Figure 2B; see also Methods).

Our initial selection for magnitude and probability coding (Eq. 9) identified also neurons that potentially coded only reward magnitude (96 of 622 neurons = 15%; Monkey A: 20 / 149; Monkey B: 76 / 473), only reward probability (76 of 622 neurons = 10%; Monkey A: 28 / 149; Monkey B: 48 / 473), or none of these variables (272 of 622 neurons = 44%; Monkey A: 71 / 149; Monkey B: 201 / 473). Thus, the neuronal sample did not include neurons with non-monotonic response functions; their responses followed inverted-U functions on reward magnitude and/or probability and could be fitted to varying degrees to Gaussian functions. They resembled previously described OFC responses that increased with the statistical variance of reward probability distributions (O’Neill and Schultz, 2010).

### Subjective economic value measure in individual neurons

Via the same axiomatic principle used to characterize behavior, we defined a neuronal value measure as the indifference point computed from the neuronal activity. This neuronal indifference point (IPn) corresponded to the probability *p*_A_ at which the neuron responded equally strongly to the two choice options; the test was repeated for different A and B magnitude levels (Figure 2A, B), as in the behavioral measures. Each option-set {A,B,C} defined one test for computing the IPn. A neuron encoding subjective economic value would respond equally to the two choice options at the IPb. The same neuron, when presented with an AC option preferred (or not preferred) to the B option, would produce a stronger (or weaker) activation. By comparing the IPn to the IPb in each test we evaluated how well a neuron encoded the behaviorally estimated subjective economic value.

To compute the IPn for each option set in each neuron coding both reward magnitude and probability, we first decoded preferences from the neuronal activity using a specifically designed choice-probability decoder (see Methods). We decoded the behavioral preference (i.e., the probability of choosing option AC over B) from the probability that the neuronal response to AC was higher than the response to B (Eq. 11, see Methods and Figure S4). We then fitted a softmax function (Eq. 3) to the decoded preferences, identifying the IPn as the *p*_A_ corresponding to a softmax value of 0.5 (Figure S4C). The IPn values (Figure 2D, black line) varied across neurons, generally differing from the corresponding behavioral IPb values (Figure 2D, green line). In some neurons (example neuron 1, Figure 2), the IPn was close to the IPb in most tests; in other neurons the IPn was larger (example neuron 2) or smaller (example neuron 3) than the IPb, in all tests. Similar characteristics in subjective value coding were also observed during binary choice (Figure S3B).

In economic terms, neurons with IPn larger than the EV-based indifference point (IPev, i.e., the *p*_A_ for which the AC and B options had equal EV) predicted a risk averse behavior, while neurons with IPn < IPev predicted a risk seeking behavior (Figure 3A). Across the population of recorded neurons, the IPn varied continuously within each test, showing heterogeneity in the coding of subjective value and a general non-conformity of the neuronal value measure with the behaviorally defined one (Figure 3B).

**Figure 3.**
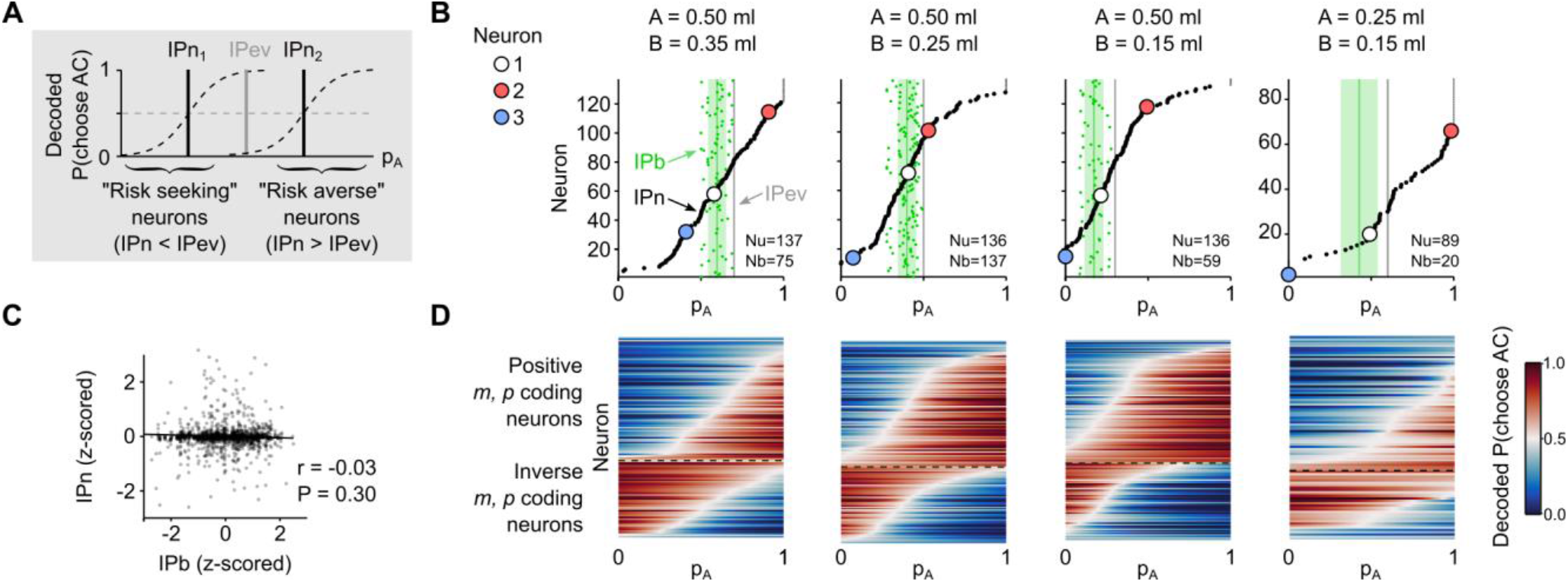
Heterogeneous coding of subjective economic value in the neuronal population. (A) Economic interpretation of diverse value codes. Neurons can be classified as encoding a risk seeking behavior or a risk averse one, based on their IPn (i.e., subjective economic value measure) being smaller or larger than the EV-based indifference point (IPev). (B) IPn measure across all recorded neurons in each of the four example tests. Each dot corresponds to the IPn coded by one neuron in one specific option-set (neurons were sorted by IPn, separately for each test). The broad and continuous variation of prospective subjective economic value measures in single neurons corresponded to the coding of different risk attitudes. Color-filled circles: example neurons 1, 2 and 3 (same as in Figure 2). The number of neurons (Nu) and behavioral sessions (Nb) varied as a subset of tests was performed in every recording session. (C) Variability of neuronal and behavioral value measures across sessions (Monkey B). The non-significant correlation between the z-scored IPn and IPb values showed that the variability in the neuronal value measure unlikely depended on the variability in the behavior. (D) Positive and inverse *m, p* coding neurons. Both positive and inverse coding neurons showed a continuous variation in IPn values in all tests. Colors (from blue to red) represent the neuronal decoded P(Choose AC) for every neuron, sorted by IPn. White color correspond to the IPn (P(Choose AC) = 0.5), red for the activity being higher for the AC option, blue for the activity being higher for the B option. Data in (A) to (D) are from Monkey B. For neuronal responses in choice trials see Figure S6.

As neurons were recorded in different daily sessions and the IPb’s varied slightly across sessions, some of the observed IPn variation could in principle be attributed to IPb variation. To test this hypothesis, we correlated z-score-normalized IPn and IPb values across sessions. The correlation coefficients between IPn and IPb values were insignificant (Figure 3C; Monkey A: Pearson’s r = −0.02, P = 0.56; Monkey B: r = −0.03, P = 0.30). Thus, the variability of IPn’s across neurons was unlikely due to variation of IPb’s across sessions. The neuronal-behavioral correlations were also insignificant when using a particular risk measure alpha that had resulted in significant correlations in a previous study (Raghuraman and Padoa-Schioppa, 2014) (Pearson’s rho from −0.01 to +0.02; P > 0.85) (Figure S5), although the differences in measures (alpha vs IP’s) and task design make comparisons difficult (see Discussion). The observed value coding variability across neurons was found in both positive and inverse *m*,*p* coding neurons (Figure 3D). Similar subjective value coding was also observed during binary choice (Figure S6).

These results highlight the heterogeneous coding of subjective value in individual OFC neurons, with the neuronal value measure continuously varying across neurons. We identified “risk seeking” neurons and “risk averse” neurons based on their specific subjective value code.

### Comparison of neuronal and behavioral subjective value measures

To quantify the correspondence between the neuronal and the behavioral subjective value measures across tests we followed two complementary approaches. First, we tested whether the neuronal activity corresponded to a noisy representation of behavior. Second, we directly compared the behavioral preferences with the preferences decoded from individual neurons.

A neuron coding the economic value would respond equally to two choice options at behavioral indifference. We counted the number of neurons with significantly different activity (t-test, P < 0.05) in response to two equally preferred options and compared it to the expected false discovery rate (FDR, 5%) (see Methods). Among the neurons recorded in four tests (17 in Monkey A, 137 in Monkey B), we found 20.6% (Monkey A) and 19.5% (Monkey B) of neurons with activity significantly different between the two options in one of the tests (randomly selected test, procedure repeated 20,000 times), a significantly higher percentage than the 5% chance level (t-test, P < 0.05). When randomly selecting gradually more tests, we found an increasing number of neurons with different activity between the two equally preferred options in at least one test; the percentages of neurons stayed above the respective chance level (Figure 4A). Thus, the number of neurons with responses deviating from behavior was higher than what would be expected from neuronal noise.

**Figure 4.**
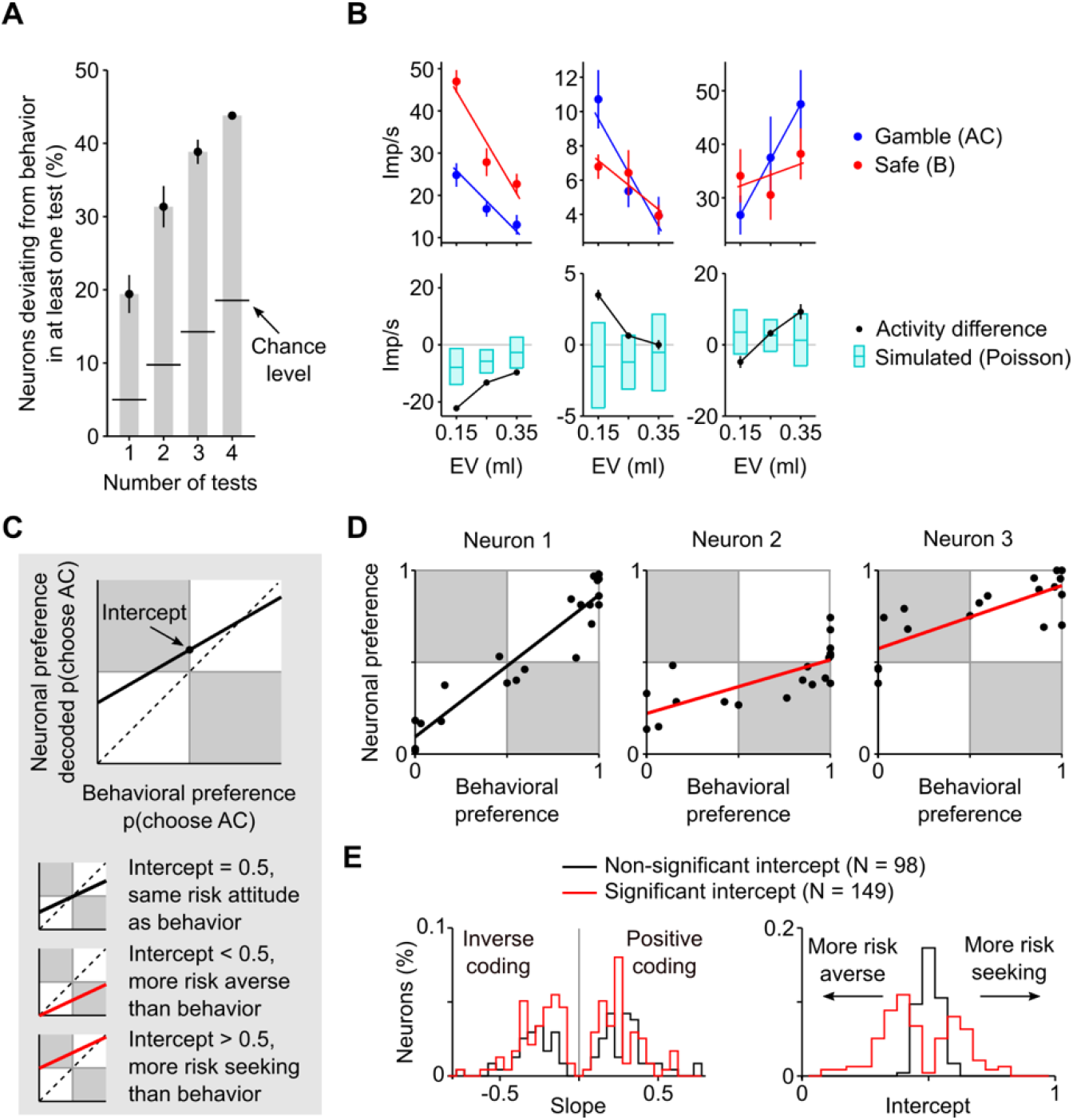
Neuronal value coding deviating from behaviorally assessed subjective value. (A) Percentage of neurons whose activity significantly differed between two equally behaviorally preferred options in at least one test (error bars: 95% confidence interval (CI)). The chance level (false discovery rate) was computed as 1 – (1 – α)^N^, where α = 0.05 (significance level), N the number of tests performed. Data from Monkey B. (B) Three simultaneously recorded neurons showing activity incompatible with the noisy (Poisson-distributed) coding of behavior. Top: neuronal response (mean ± SEM) to gambles with m = 0.5 ml (blue) or safe (red) options. Bottom: difference in response rate between gamble and safe options with the same EV, in the recorded neuron (black) and in the corresponding (same activity range) simulated neuron (cyan; area: 95% CI). The three neurons were recorded in the same session (left neuron form electrode 1, remaining two neurons from electrode 2), ensuring that response differences were not due to differences in behavior. (C) Decoding the risk attitudes from the neuronal activity. Top: the neuronal preferences (PN, decoded based on the overlap of activity distributions between pairs of options) were regressed on the behavioral preferences P_B_ (Eq. 12). The regression slope (*a*) quantified the strength of preference coding, while the intercept (*b* + 0.5) represented the relation with the behavior. Bottom: a neuron with *b* = 0 would consistently encode the behavioral risk attitude across all tested gambles; *b* < 0 (or *b* > 0) would correspond to a more risk averse encoded attitude (or a more risk seeking one) than behavior. Behavioral preference is defined as the probability of choosing option AC over option B; neural preference is defined as percentage of trials with higher activity for option AC than for option B. (D) Example neurons encoding preferences compatible with behavior (neuron 1) or incompatible with behavior (neurons 2, 3) (same 3 neurons as in Figure 2C, D). Each dot represents the relation between neuronal and behavioral preference measures for one pair of options {B, AC}. The white regions represent areas of decoded preferences that correctly predicted behavior, while dots in the gray regions predicted a preference that was opposite to the behavioral one. The top-left number is the percentage of correct predictions. (E) Distribution of neuronal response types (Monkey B). Left: both positive and inverse *m, p* coding neurons (*a* > 0 and *a* < 0, respectively) showed a significant (red) or non-significant intercept (black). Right: distribution of intercept values. For the positive coding neurons, an intercept > 0.5 (*b* > 0; t-test, P < 0.05) corresponded to more risk seeking preferences compared to behavior, while an intercept < 0.5 corresponded to more risk averse ones across all tested gambles.

We then investigated whether the recorded activity was compatible with the noisy coding of behavior, away from the indifference point. As the cortical activity is often assumed to be Poisson-distributed, we simulated the activity of neurons coding the economic value assuming Poisson-distributed activity rates (see Methods). The activity of value-coding neurons was expected to fall within the 95% confidence interval (CI) of the simulated activity. We found percentages of neurons with activity falling outside the simulated CI that were higher than the expected 5% FDR. In particular, when considering three pairs of options (gamble vs safe) with EV equal to 0.15 ml, 0.25 ml and 0.35 ml, we respectively found 33%, 29% and 23% of neurons that deviated from the simulated activity. Three example neurons, recorded simultaneously, responded differently from the simulated behavior-compatible neurons, in opposite directions (Figure 4B); this example demonstrated the significantly different value-coding pattern across neurons, and confirmed that the results were not due to an imprecise (i.e., noisy) evaluation of behavior.

To characterize the relation between neuronal and behavioral economic value, we directly compared the neuronal decoded preferences (as used above for computing the IPn’s) with the behavioral choice preferences, across all tested option-pairs. Rather than selecting neurons according to their magnitude and probability coding, as used for single-neuron examples (Figures 2 and 3; Eq. 9), we tested OFC neurons that showed significant correlations between neuronal preferences P_N_ and behavioral preferences P_B_ across all tests (Eq. 12). This procedure allowed us to distinguish between activities that were compatible with the behavior and incompatible activities. The regression slope identified the strength of preference coding, while the intercept represented the relation with the behavioral risk attitude. An intercept of 0.5 (defined at the x-axis value of 0.5, see scheme of Figure 4C) corresponded to a neuron coding subjective value compatibly with behavior; different intercepts identified neurons encoding a more risk seeking attitude (intercept > 0.5) or a more risk avoiding one (intercept < 0.5), as compared to behavior.

When applying this selection criterion of correlated neuronal and behavioral preferences (P_N_ and P_B_; Eq. 12), we found OFC neurons that encoded preferences compatible with behavior (Figure 4D, example neuron 1) or incompatible with behavior (example neurons 2, 3), thus demonstrating neuron-specific risk attitudes across all tests. When applying the selection criterion to all 622 task-responsive neurons, we found 295 neurons (47%) that showed a significant slope coefficient (Monkey A: 48 neurons, Monkey B: 247 neurons) (P < 0.05, positive or negative regression slope coefficient, Eq. 12), demonstrating a degree of preference coding (i.e., significant correlation between neuronal and behavioral preferences) (Figure 4E, left). Of these 295 neurons, 118 neurons (40%) had an intercept between the neuronal and behavioral preferences that was insignificantly different from 0.5 (Monkey A: 20 neurons, Monkey B: 98 neurons) (P > 0.05), suggesting that their preference code was compatible with behavior; the remaining 177 neurons (60%) had an intercept significantly different from 0.5 (Monkey A: 28 neurons, Monkey B: 149 neurons) (P < 0.05, regression intercept coefficient), resulting in neurons coding a different risk attitude compared to behavior (Figure 4E, right). When we applied the stricter statistical threshold of P < 0.01, we found 213 neurons among the 622 task-responsive neurons (34%) that showed a significant positive or negative regression slope coefficient (Eq. 12; Monkey A: 35 neurons, Monkey B: 178 neurons), demonstrating a degree of preference coding (i.e., significant correlation between neuronal and behavioral preferences). Of these 213 neurons, 105 neurons (49%) had an intercept between the neuronal and behavioral preferences insignificantly different from 0.5 (Monkey A: 17 neurons, Monkey B: 88 neurons) (P > 0.01), suggesting a preference code compatible with behavior; the remaining 108 neurons (51%) had an intercept significantly different from 0.5 (Monkey A: 18 neurons, Monkey B: 90 neurons) (P < 0.01, regression intercept coefficient), suggesting neurons coding a different risk attitude compared to behavior. Taken together, this analysis quantified the correspondence between the neuronal and the behavioral preference measures, showing the existence of distinct subpopulations of neurons with different relations to the behavioral risk attitudes.

These results show that the heterogeneous responses of OFC neurons robustly reflected a multitude of choice behaviors, continuously varying from risk seeking to risk aversion, including a subpopulation of neurons with behavior-compatible risk attitudes.

### Heterogeneous signals for utility and probability weighting in individual neurons

To investigate the source of the observed heterogeneity in the coding of the IPn (i.e., the subjective economic value) across OFC neurons, we analyzed the Cue responses to the full range of reward magnitude and probability levels required for testing the continuity axiom. When comparing the responses to safe rewards with the responses to gambles, two features emerged (Figure 5A): (1) the responses were a nonlinear function of the EV, and (2) the relative sensitivity to magnitude (red) and probability (blue) varied across neurons. These two characteristics defined the “risk attitude” of individual neurons at each EV level: a higher activity for gambles compared to EV-matched safe options demonstrated the coding of a risk seeking attitude (Fig 5A, example neurons 1, 3), while a higher activity for the safe compared to the gamble option identified the coding of a risk averse attitude (Fig 5A, example neuron 2). Similar heterogeneous neuronal coding of risk attitude was found with the 64 chosen value neurons recorded in choice trials. Example neurons from both animals are shown in Figure S7A.

**Figure 5.**
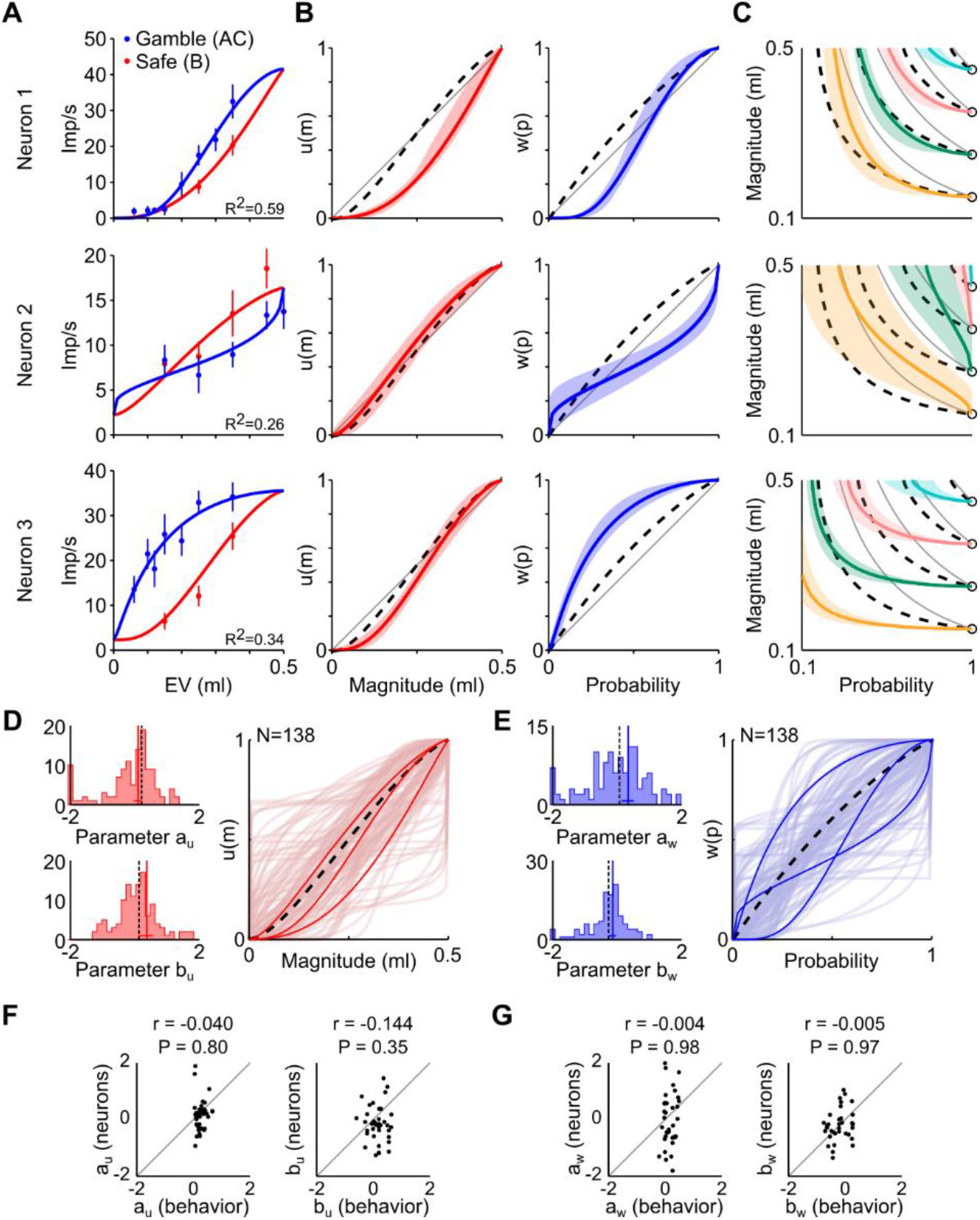
Heterogeneous neuronal economic functions across single neurons. (A) Single neurons’ coding of magnitude and probability. Activity (mean ± SEM; Imp/s: impulses per second) of three example neurons (same 3 neurons as in Figures 2C, D and 4D) to different reward magnitudes (red) and probabilities (blue). The curves resulted from the subjective economic value function *v(m,p)* directly fitted to the activity of individual neurons (only one set of magnitude and probability levels are shown (red: *p* = 1, blue: *m* = 0.5), while the fit was performed on all tested *m* and *p* levels). For neuronal responses in choice trials see Figure S7. (B) Neuronal and behavioral utility and probability weighting functions. The functions computed for each neuron (continuous lines) varied across neurons and differed from the behavioral ones (dashed lines). Gray lines: objective values (linear functions). Shadows show standard deviations. (C) Neuronal and behavioral indifference map. Comparison between neuronal ICs (colored lines), behavioral ICs (dashed lines) and EV-curves (gray lines). Shadows show standard deviations; 95% confidence intervals (Figure S9) confirmed significant differences between neuronal and behavioral data. (D), (E) Distribution of best-fitting neuronal economic functions’ parameters (left) and corresponding functions (right) across neurons. Data from Monkey B. (F), (G) Variability of neuronal and behavioral economic functions’ parameters. The parameters of both utility (F) and probability weighting (G) functions were not significantly correlated between neurons and behavior, suggested that the variation in neuronal economic functions was not due to behavioral variations across sessions.

The encoded risk attitude thus depended on the relative strength of the activity for the safe option (red) compared to that for the gamble option (blue), at each EV. Diverse nonlinear responses produced varied risk attitudes throughout the range of rewards; similar data was obtained for both positive and inverse *m, p* coding neurons, in both animals (Figure S8).

We linked this idea with the economic constructs of utility and probability weighting functions, which correspond to subjective nonlinearities in the evaluation of reward magnitudes and probabilities, respectively. To this aim, we directly fitted a nonlinear economic model to the neuronal activity. In brief, we defined the subjective economic value *v(m,p)* of a gamble through the multiplicative subjective economic value model, using separate 2-parameter Prelec functions for the nonlinear contributions of magnitude (*u*) and probability (*w*) (Eqs. 4 and 5, see Methods). Then we fitted the value function *v(m,p)* to the neuronal activity rate *N* via nonlinear least squares (Eq. 13). We thus characterized the activity of each neuron across all tested gambles with a set of 6 parameters (for the 2-parameter utility function, the 2-parameter probability weighting function, and 2 free parameters for the least squares fit). This fit identified an overall neuronal value function linking the nonlinear neuronal utility function and the nonlinear neuronal probability weighting function.

The recovered neuronal functions highlighted the neurons’ heterogeneity in relation to behavior, showing specific nonlinearities in the coding of magnitudes and probabilities (Figure 5B). Using the neuronal economic functions, we reconstructed the ICs encoded by individual neurons: the neuronal ICs (Figure 5C). One example neuron (Figure 5C, top) produced ICs close to the behavioral ones, although its *u* and *w* functions differed from the behaviorally elicited ones (Figure 5B); other neurons produced ICs that were clearly incompatible with the behavior, being warped toward the right (less risk seeking than the choices; Figure 5C, middle) or to the left (more risk seeking than the choices; Figure 5C, bottom), although their utility functions were similar to the behavioral ones (Figure 5B). In example neurons 2 and 3, the main contribution to the coded risk attitude came from the nonlinearity in the coding of probabilities, while in neuron 1, the combination of nonlinear *u* and *w* contributed a subjective economic value code compatible with behavior. Similar heterogeneous economic functions and ICs across different OFC neurons were found with the 64 chosen value neurons (Figure S7B, C).

Across the neuronal population, the neuronal *u* and *w* functions generally differed from the behavioral ones (Figure 5D, E). As the behavioral economic functions also varied across sessions, reflecting fluctuations in the choice patterns (Figure S10), we tested the hypothesis that the neurons were simply following the behavioral variations. We found non-significant correlation coefficients between the neuronal and behavioral parameters of both the utility and probability weighting functions (Figure 5F, G), suggesting that the variation in neuronal economic functions was unlikely due to behavioral variations across different sessions. Neuronal-behavioral correlations were also insignificant (Pearson’s rho from −0.03 to +0.10; P > 0.32) when using a risk measure derived from power function parameters for utility and probability weighting (Figure S11), thus replicating our findings with the alpha risk measure used by Raghuraman and Padoa-Schioppa (2014) (Figure S5) (see Discussion).

These data show that the diverse coding of subjective economic value across single OFC neurons reflected the heterogeneous nonlinearity in the coding of reward magnitude and probability. These characteristics were captured by the neuronal economic utility and probability weighting functions recovered in single neurons, which varied greatly across neurons, as compared to the behavioral ones.

### A reliable population code for subjective economic value

Our axiom-based characterization of the activity of individual neurons allowed us to identify neuron-specific utility and probability weighting functions. However, their individual activity did not generally correspond to the monkey’s choice behavior. This puzzling result opens a crucial question about the role of OFC neurons in choice: might behavior-compatible subjective economic values be represented at a population level?

To test how a neuronal population signal would correlate with the animal’s behavior, we tested the 295 OFC neurons that had been selected based on their correlated neuronal and behavioral preferences (P_N_ and P_B_; Eq. 12). To do so, we combined the recovered economic function parameters of different subpopulations of neurons and compared them with the behavioral ones.

We selected three neuronal populations based on the analysis relating behavioral and neuronal decoded preferences (Figure 4C). The first population included the neurons whose decoded preferences were compatible with the behavior (i.e., neurons for which the intercept of the regression line between neuronal and behavioral preferences was close to 0.5). We refer to these as the behavior-compatible neurons. The other two populations included neurons encoding preferences that were more risk averse (intercept < 0.5) or more risk seeking (intercept > 0.5) than the behavior (behavior-incompatible neurons). We decoded preferences at the population level following the heuristic argument that, in noisy multi-component systems, accuracy can be improved by computing the output based on the majority of single-component outputs (von Neumann, 1956). We thus computed the preference (i.e., the proportion of gamble choices in a gamble-safe choice pair) as the percentage of neurons encoding a preference for the gamble option (see Methods).

The unweighted (unbiased) combination of behavior-compatible neurons into a neuronal subpopulation showed that the neuronal preferences correlated strongly with the behavioral preferences (Monkey A: Pearson’s r = 0.85, P = 6.1e-10; Monkey B: r = 0.93; P = 6.2e-14), correctly predicting most behavioral choices (Monkey A: 78%, Monkey B: 87%) (Figure 6A). The behavior-compatible neurons showed utility and probability weighting functions that matched well the corresponding behavioral utility and probability weighting functions; the behavioral functions fell within the 95% confidence intervals of the neuronal functions (Figure 6B). We calculated the population-derived economic functions’ parameters as the median of the parameters obtained in individual neurons and estimated the functions’ 95% CI through a bootstrap procedure (see Methods). Medians were deemed more appropriate than means, as medians control for the asymmetry in the parameters’ distributions and the contribution of outliers. The goodness of the behavioral prediction was confirmed by the precise overlap between neuronal and behavioral ICs, which fell within the 95% confidence intervals of the neuronal ones (Figure 6C).

**Figure 6.**
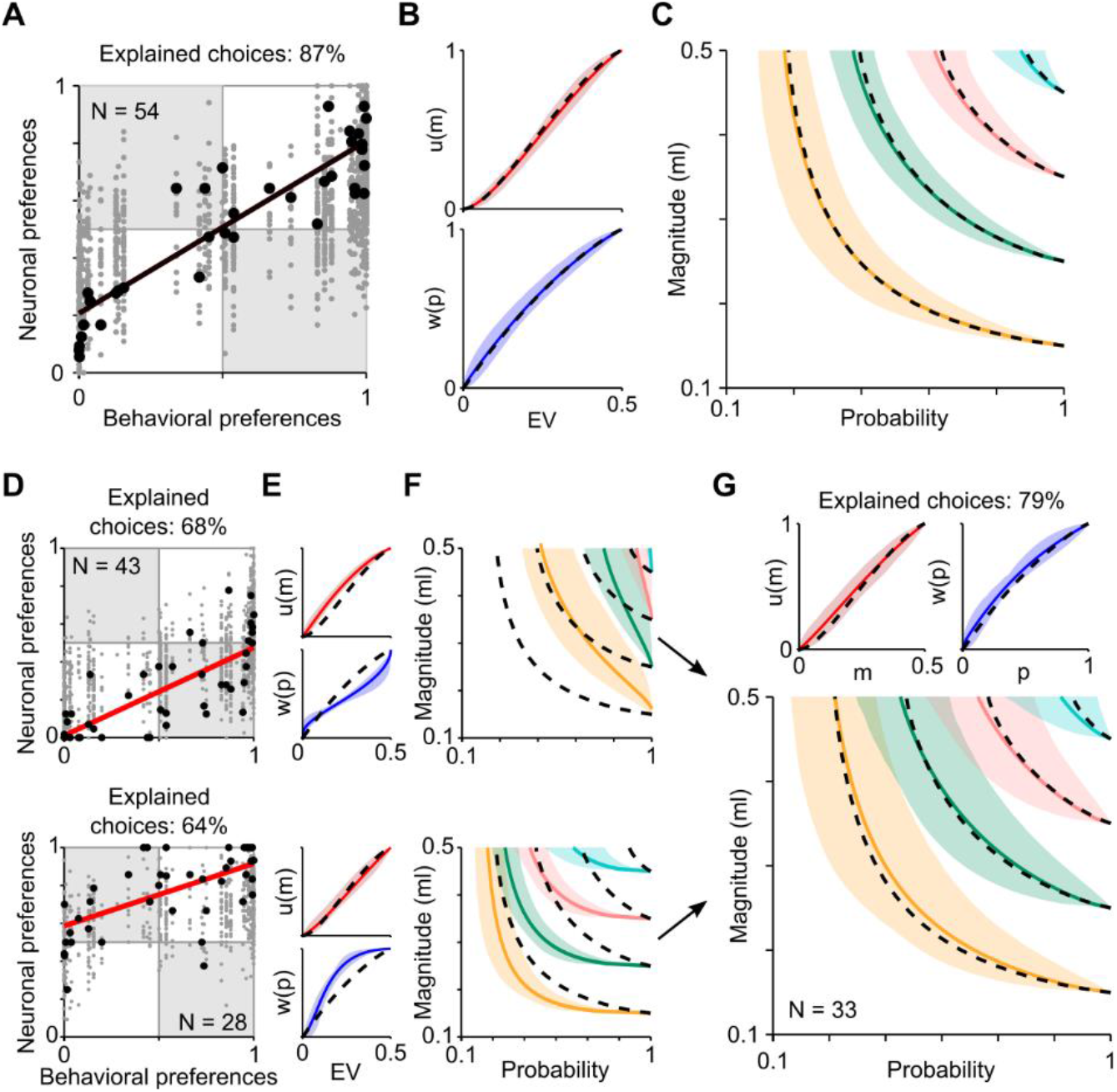
Reliable population code of subjective economic value. (A) Population-decoded preferences for the behavior-compatible neurons (N = 54). Each black dot represents the relation between behavioral and neuronal-population preference. Preference (i.e., the probability of choosing the gamble AC) was computed from the neuronal population as the percentage of neurons encoding a preference for the gamble option (gray dots: preferences decoded from each neuron). (B) Population-encoded economic functions. Utility (u, red) and probability weighting (w, blue) functions obtained from the median parameters across all behavior-compatible neurons. The 95% CI (colored area) computed on the neuronal economic functions included the behavioral curves (black dashed lines). (C) Indifference curves (ICs). The neuronal ICs, computed from the economic model’s parameters (colored lines and 95% CI regions), matched the behavioral ones (black dashed lines) in the behavioral-preference neuronal population (N = 54 neurons). (D), (E), (F) Neuronal (decoded) and behavioral preferences (D), economic functions (E) and ICs (F) for the behavior-incompatible neurons (top: neurons more risk averse than behavior; N = 43) (bottom: neurons more risk seeking than behavior; N = 28). (G) A combination of neurons from the behavior-incompatible subpopulations encoded behavior-compatible economic functions (top) and ICs (bottom) (N = 33 neurons). For neurons’ selection method see Figure S12. Data in (A) to (G) are from positive *m, p* coding neurons in Monkey B. For inverse coding neurons of Monkey B and for positive coding neurons of Monkey A, see Figure S14.

Then we tested unweighted combinations of the two behavior-incompatible neuron types into two respective neuronal subpopulations. In the more risk averse neuronal subpopulation (Figure 6D, top) the neuronal and behavioral preferences were significantly correlated (Monkey A: Pearson’s r = 0.74; P = 1.6e-6; Monkey B: r = 0.75; P = 1.2e-6); in the more risk seeking subpopulation (Figure 6D, bottom), correlation was significant only for Monkey B (Monkey A: Pearson’s r = 0.18; P = 3.3e-1; Monkey B: r = 0.68; P = 1.7e-6). As expected, these two subpopulations explained a lower proportion of choices compared with the behavior-compatible neurons (Monkey A: 56% and 66% for the more risk averse and for the more risk seeking populations, respectively; Monkey B: 68% and 64%). Consistently, the economic functions elicited from the two subpopulations were significantly different from the behavioral ones (behavioral curves falling outside the 95% CI of the neuronal ones). In particular, the probability weighting function was different between the two subpopulations: the more risk averse neuronal population underweighted the high probabilities (Figure 6E, top), while the more risk seeking population overweighted them (Figure 6E, bottom). This resulted in ICs warped toward the right (confirming a significantly risk averse encoded attitude) or toward the left (risk seeking encoded attitude), with 95% CIs not comprising the behavioral curves (Figure 6F).

The question arose whether the unreliable neurons might contribute to a reliable value computation. Therefore we tested how well the two behavior-incompatible subpopulations together might match the animal’s behavior. We analyzed only neurons whose responses showed an intercept between neuronal and behavioral preferences that differed significantly from 0.5 (see Figures 4C, D and 6D); we defined the sum of squared errors (SSE) along the probability dimension as metric for the difference between neuronal ICs and behavioral ICs. We then looked for a subpopulation of neurons that most closely matched the behavioral ICs (i.e., with the lowest SSE). This subpopulation was identified by first sorting the neurons based on the absolute value of their intercept and then computing the SSE for subpopulations of increasing numbers of neurons. The SSE reached a minimum value for N = 33 neurons (Figure S12), which indicated the subpopulation of neurons that produced the best fit to behavior; adding more neurons produced less reliable ICs. Analysis with this population produced neuronal economic functions and ICs that overlapped well with the behavioral ones (Figure 6G). These results were also obtained with a stricter statistical threshold of P < 0.01 (Figure S13). Thus, the unweighted (unbiased) combinations of unreliable neurons resulted in good correlations with the animal’s behavior, even though the individual neurons coded reward value in a heterogeneous manner that reflected opposite behavioral patterns.

These results, obtained from Monkey B’s positive *m, p* coding neurons, were replicated in the population of inverse coding neurons (Figure S14A). Data from Monkey A, although noisier (given the lower number of available neurons), showed the same pattern (Figure S14B).

In addition to the different population responses of the behavior-compatible and the behavior-incompatible OFC neurons shown in Figure 6, we pooled the responses of these distinct populations into a single common neuronal population code. The common neuronal population preferences correlated well with the behavioral preferences and predicted the behavioral choices to 87% correct (Figure 7A). The utility and probability weighting functions matched well the corresponding behavioral functions (Figure 7B). The neuronal and behavioral ICs overlapped well; the behavioral ICs fell within the 95% confidence intervals of the neuronal ICs (Figure 7C). These data demonstrate a reliable representation of the behaviorally assessed subjective economic values in a mixed population of reliable and unreliable OFC neurons.

**Figure 7.**
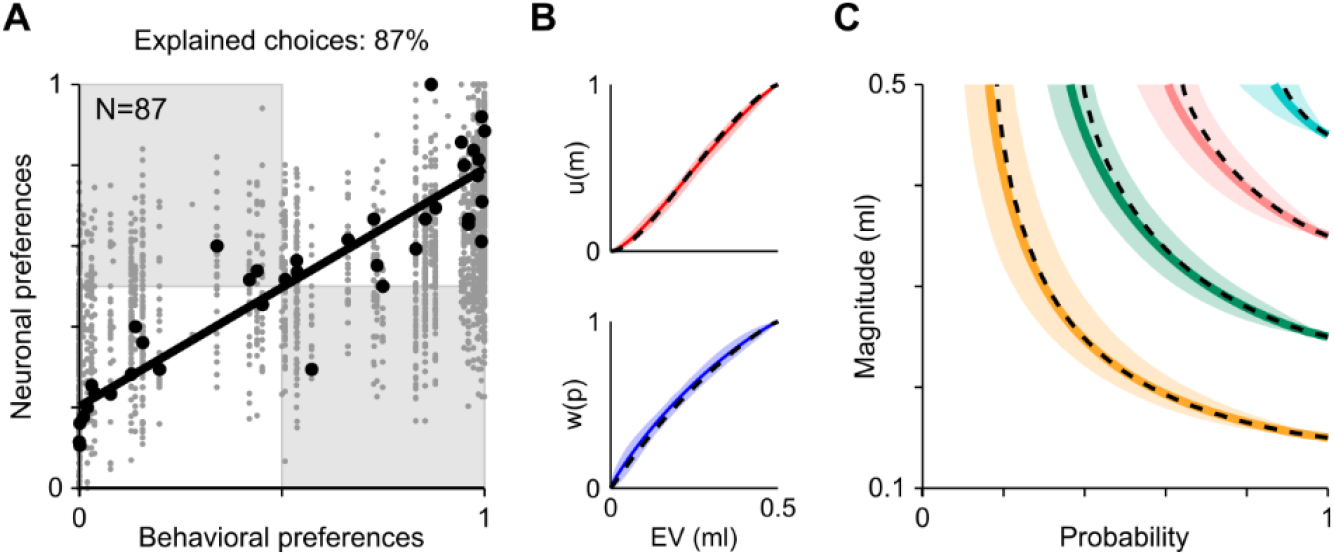
Reliable coding of economic value by a mixed population of reliable and unreliable neurons. (A) Correlation between behavioral and neuronal preferences in a population of behavior-compatible neurons (N = 54) and behavior-incompatible neurons (N = 33 neurons; total of 87 neurons). Conventions as in Figure 6. (B) Closely matching utility function u(m) and probability weighting function w(p) between behavior (dashed lines) and neurons (solid lines with 95% confidence intervals). (C) Closely matching indifference curves between behavior and neurons.

Overall, these data show that subpopulations of heterogeneous OFC neurons encoded distinct subjective economic values corresponding to different behavioral attitudes. Our population analysis revealed that the combination of unreliable responses of individual neurons resulted in a reliable population code for subjective value.

## DISCUSSION

Using extensive and structured variations of reward magnitude and probability guided by the continuity axiom of EUT, we found that the population activity of OFC reward neurons resulted in a neuronal code that closely matched the behavioral choices. Thus, the unreliable responses of individual OFC neurons translated into a population signal that reliably coded subjective reward value composed of utility and weighted probability. Thus, the reliable OFC population signal constitutes an emergent property that provides a non-trivial advance over the known reward signals of individual OFC neurons. The advance lies in the fact that unbiased combinations of unreliable individual reward signals do not necessarily result in population signals that reliably correlate with behavioral choices.

In the experiment, we used the previously defined subjective economic value measure that complied with the continuity axiom of EUT (Figure 1). Axiom compliance demonstrated understanding of the employed choice options and meaningful choice behavior, which constitutes a necessary condition for assessing the reliability of neuronal signals for subjective reward value defined by reward magnitude and probability. The value inferred from behavioral choices reflected nonlinear utility and probability weighting functions (Ferrari-Toniolo et al., 2021), as conceptualized by PT (Kahneman and Tversky, 1979), whereas the mean-variance approach had provided inferior behavioral fits (Ferrari-Toniolo et al., 2021). We analyzed all OFC neurons whose Cue responses varied significantly with both reward magnitude and reward probability. Despite the stringent axiomatic approach and the preselection of neurons, the value code of OFC neurons was heterogeneous and failed to correspond to the subjective economic value inferred from the observed behavioral choices (Figure 2). Individual neurons encoded diverse risk attitudes (Figures 3 and 4). The diverse coding applied to both utility and probability weighting (Figure 5). These heterogeneous responses occurred in both zero-alternative trials with minimal confounds from unchosen options and in choice trials with more direct behavioral-neuronal correspondence. However, the simple combination of unreliable responses of individual neurons resulted in a reliable population code that corresponded to the monkeys’ risky choices with high precision (Figure 6). Thus, OFC neurons encoded subjective economic value as a population of heterogeneous individual elements.

### Design

The continuity axiom of EUT, central in defining a numerical measure of subjective value, guided our structured tests of a large variety of reward magnitudes and probabilities that resulted in the surprising finding of unreliable value coding in individual OFC neurons. In principle, compliance with the axiom must be verified for any possible set of gambles. To assess subjective value from choices according to the axiom, we varied reward magnitudes and probabilities across the whole experimental space as much as was practically possible. This large variation of testing parameters allowed us to characterize the responses of single neurons in subjective economic value terms, thus revealing the striking heterogeneity of value coding across neurons as well as across reward levels. By directly fitting the economic model to the neuronal data, we captured the complex pattern of value measures of individual neurons across the whole magnitude and probability space. We defined subjective economic value as the product of the underlying economic functions (utility and probability weighting) and used the model for both the behavioral choices and the neuronal responses. Assuming that single neurons encoded subjective economic value, the procedure identified separately in each neuron the best-fitting utility and probability weighting functions defining that value. We thus characterized the variation of activity patterns across neurons in terms of their value-generating economic functions.

Utility is often modeled as a power function in the gain domain (Yamada et al., 2013; Chen and Stuphorn, 2018). However, PT postulates a canonical inflected utility function that is convex for reward magnitudes below a reference point and concave above the reference point (Kahneman and Tversky, 1979). To capture a broad range of utility shapes reflecting a wide range of risk attitudes irrespective of gain or loss domain, we employed the two-parameter Prelec function that allows for convex, concave, or mixed utility functions (regular or inverse S-shape) that are valid for the studied gain domain. The choice of an inflected utility function is particularly important for two aspects: first, previous studies have shown that monkeys are risk seeking for small reward magnitudes, while being risk averse for larger rewards, resulting in an S-shaped utility function (Stauffer et al., 2014; Bujold et al., 2021; Ferrari-Toniolo et al., 2021), which explains choices better than a power-shaped one (Ferrari-Toniolo et al., 2021); second, for neuronal response, the Prelec function is ideal for flexibly characterizing the nonlinear coding of reward variables, without biasing the results by imposing a pre-determined profile.

### Comparison with correlations using different risk measures

In contrast to our insignificant correlations between neuronal and behavioral measures (Figures 3C, 5F, G), a previous study reported neuronal-behavioral correlations that were indeed significant (Raghuraman & Padoa-Schioppa 2014). Our correlations remained even insignificant when using the same ‘alpha’ risk measure as in the previous study or when using a risk measure derived from our power functions for utility and probability-weighting (Figure S5). Thus, the different measures used between the previous study and our current study unlikely explain the discrepancies, and we should consider the differences in the selection of neurons and some task details, as follows: Selection of neurons: due to the different task design, our stimuli indicating the two choice options were not unambiguously distinct and thus difficult to identify by the animal. Hence, we could not identify object value neurons that code only the value of one specific choice object (or ‘offer value’ neurons according to Padoa-Schioppa and Assad, 2006). By contrast, both the previous study and our current study could identify chosen value neurons that code the value of the option the animal is going to choose or has just chosen. Thus, the neuronal sample differed between the two studies being compared.

Task details: the tasks differed in terms of trial sequence (delay between stimulus presentation and choice with Raghuraman & Padoa-Schioppa (2014), reaction time task in our study) and in terms of larger number of probability levels tested in our study (while the tested magnitude levels were about similar).

### Meaningful value signal from heterogeneous neurons

Our results highlight the nonlinear coding of reward magnitude and probability in individual OFC neurons. By fitting a subjective economic value model to the neuronal activity, we related the observed neuronal nonlinearities to the subjective economic value functions inferred from the behavioral choices. This link allowed us to compare neuronal activity to choice behavior through the best-fitting economic functions in the two domains. According to previous studies, monkeys’ choices indicate nonlinear weighting of both reward magnitude and probability (Stauffer et al., 2014; Bujold et al., 2021; Ferrari-Toniolo et al., 2021). Thus, we hypothesized that neuronal coding of subjective economic value should also be nonlinear for magnitudes and probabilities and should correspond to the nonlinear functions inferred from behavioral choices.

Our analysis was restricted to OFC neurons whose Cue responses varied significantly with both reward magnitude and reward probability. Despite this well-defined preselection, the estimated economic function parameters of neuronal responses were heterogeneous across individual neurons and amounted to a rather wide distribution. This diversity was true for both utility and probability weighting functions. Risk attitudes can be reflected in the relative shapes of the two economic functions: risk seeking, for instance, can be represented by a convex utility function as well as by a concave probability weighting function. More complex functions’ shapes (e.g., S or inverse-S) can reflect more complex patterns of risk attitude that depend on the specific gambles being offered. Thus, the diversity of economic function parameters recovered from individual OFC neurons indicated a broad range of risk attitudes. The different neuronal subpopulations reflected risk seeking, risk aversion or mixed risk attitudes.

Previous studies on OFC neurons estimated subjective value in particular situations, such as changing choice preferences, spontaneous satiation and temporal discounting (Tremblay and Schultz, 1999; Padoa-Schioppa and Assad, 2006; Kobayashi and Schultz, 2008; Padoa-Schioppa and Cai, 2011). By contrast, our study, using economic axiomatic theory, estimated a general form of subjective economic value derived from mathematical economic functions for utility and probability-weighting. Using this approach, we related the activity of individual OFC neurons to these value-estimating functions. This strategy was instrumental in identifying and characterizing the heterogeneous coding of subjective reward value across OFC neurons.

Unreliable coding of reward value has also been observed in dopamine neurons (Dabney et al., 2020). The identified ‘optimistic’ and ‘pessimistic’ neurons show different sensitivity at specific reward levels (i.e., varying slopes for reward prediction errors), and the full reward information is only contained in the whole neuronal distribution. This heterogeneity corresponds to the presently observed heterogeneity in nonlinear utility and probability coding in OFC neurons. These two types of neurons encode their respective reward variable with different weights, namely reward prediction errors in the case of dopamine neurons, and subjective value composed of utility and probability in the case of OFC neurons. And similar to the dopamine population response that provides better reinforcement than individual neuron response, the OFC population provides a more reliable code for economic choice than most individual OFC neurons. Thus, the emergence of reliable reward coding from unreliable individual elements may constitute a broader feature of neuronal reward signals.

In addition to utility coding, OFC neurons encode a probability weighting function that constitutes the other fundamental component of subjective economic value. Nonlinear probability weighting occurs also in the human frontal cortex and striatum, as shown by functional imaging (Tobler et al., 2008; Hsu et al., 2009). As these human imaging studies measure activity from thousands of neurons at once, their results may correspond to the currently observed OFC population signal for probability weighting in monkeys that derived from heterogeneously coding individual OFC neurons. While these results demonstrate the general existence of neuronal signals for probability weighting in the brain, our findings specify that the probability weighting signal in monkey OFC derives from a heterogeneous code in individual neurons.

### Conclusions

The heterogeneity in the coding of reward magnitude and probability across individual neurons suggests possible neuronal implementations of the value computation mechanism. Responses in a few individual neurons closely reflected the animal’s choices, but these correlations often held only for a limited number of option sets. By contrast, the responses of most OFC neurons corresponded to different choice behaviors (e.g., risk seeking or risk averse). In terms of utility function, this idea conforms with a class of economic models of stochastic choice in which a decision maker has a probability distribution of possible utility functions (Agranov and Ortoleva, 2017). In analogy, a decision-maker may also have a distribution of nonlinear probability weighting functions. In repeated choices, different utility functions and/or probability weighting functions may be employed, which at the neuronal level might correspond to different sets of OFC neurons being recruited. While further studies are required for testing such possibilities, our study may contribute a basic framework for an experimental validation of such a mechanism.

Overall, these results present an insight into brain mechanisms for the computation of subjective economic value. The population of OFC neurons encodes subjective economic value in good correspondence to choices, but most individual OFC neurons show unreliable relationships to subjective economic value. The results suggest that unreliable individual neuronal elements can achieve reliable reward processing as a population. The present populations signal was obtained by averaging signals from individual neurons; with such a population signal, the OFC may correspond to the ‘Majority Organ’ of Von Neumann (1956) that achieves correct performance from noisy input elements. The noise in OFC may consist of synaptic weight changes, trial-by-trial response changes in individual neurons, and changing neuronal subpopulations that are active in a given choice situation.

## Acknowledgements

The work was supported by a Wellcome Trust Principal, Research Fellowship to WS, by Wellcome Grants WT 095495 and WT 204811, and by European Research Council (ERC) Advanced Grant 293549. We thank Aled David and Christina Thompson for animal and technical support, Dr. Polly Taylor for expert anesthesia, Dr. Henri Bertrand for veterinary support and Dr. Federico M. Echenique and Dr. David M. Grether (Caltech) for helpful comments.

## Author contributions

Experimental design, SF-T and WS; Experimentation, SF-T; Data analysis, SF-T; Writing and Editing, SF-T and WS.

## METHODS

### Rhesus monkey (*Macaca mulatta*)

Two adult male macaque monkeys (*Macaca mulatta;* Monkey A, ‘Tigger’; Monkey B, ‘Ugo’), weighing 12.6 kg and 13.8 kg, respectively, were used in the experiments. The animals were born in captivity at the Medical Research Council’s Centre for Macaques (CFM) in the UK. Both animals had not been used in other studies; behavioral results have been published in detail (Ferrari-Toniolo et al., 2021).

All experimental procedures had been ethically reviewed and approved and were regulated and continuously supervised by the following institutions and individuals in the UK and at the University of Cambridge (UCam): the Minister of State at the UK Home Office, the Animals in Science Regulation Unit (ASRU) of the UK Home Office implementing the Animals (Scientific Procedures) Act 1986 with Amendment Regulations 2012, the UK Animals in Science Committee (ASC), the local UK Home Office Inspector, the UK National Centre for Replacement, Refinement and Reduction of Animal Experiments (NC3Rs), the UCam Animal Welfare and Ethical Review Body (AWERB), the UCam Governance and Strategy Committee, the Home Office Establishment License Holder of the UCam Biomedical Service (UBS), the UBS Director for Governance and Welfare, the UBS Named Information and Compliance Support Officer, the UBS Named Veterinary

### Experimental design

We presented each animal with choices between a gamble and a safe option. These options were mutually exclusive and collectively exhaustive and appeared simultaneously and at equal distance from the animal in front of it on a computer monitor. All gambles had one non-zero outcome, which could vary in magnitude (*m*, from 0 ml to 0.5 ml in 0.05 ml steps) and probability (*p,* from 0 to 1, in 0.02 steps) and were represented respectively by the vertical position and width of a horizonal bar (Figure 1A). A safe option represented a degenerate gamble (*p* = 1). In each option, we set the magnitude and probability of reward independently. The bar’s horizontal position was randomly shifted on each trial (within the two vertical bars) to avoid delivering information about the expected value (EV) of the gamble (a fixed bar’s edge position would completely identify the gamble’s EV).

A trial started when a cross appeared at the center of the computer monitor (eye-screen distance 50 cm); a cursor, controlled by the monkey through left-right joystick hand movements, was required to be screen-centered, and after a control period (1 – 1.5 sec) two visual Cues appeared, representing the two choice options, whose fixed left and right positions alternated pseudorandomly. We inferred subjective reward value from observable choice between two options. Thus, the animal revealed its preference at the time of choice. Such ‘revealed preferences’ contrast with ‘stated preferences’ that are expressed, typically in humans, at some interval or distance from the actual choice and therefore reflect subjective value in a less reliable manner. In our experiment, the monkey revealed its preference by moving the cursor toward the chosen option. After holding it there for 1 sec, a liquid reward (water) was delivered through a spout directly in front of the animal’s mouth. The reward was contingent on the selected option’s magnitude and probability (Figure 1B). Trials containing options with different reward magnitude and probability levels were pseudorandomly interleaved.

We estimated subjective economic value at choice indifference against a fixed reward. A reward was assumed to have the same subjective value as its alternative when both rewards were chosen with equal probability (*P* = 0.5 each of two rewards in binary choice). To do so, we held reward magnitude of one option constant (option B with reward probability *p* = 1.0; Figure 1C) and varied the reward probability of the alternative option across the full probability range of 0 and 1 (option AC, with reward magnitude of component C fixed at *m* = 0). We estimated choice indifference from an S-shaped psychophysical choice function fitted to the probabilities of choosing option A over option B (Figure 1C, D; logistic regression, see Eq. 6 below). We plotted twodimensional indifference points (IP) by positioning each equally frequently chosen option with its specific reward magnitude and probability at the intersection of the x-coordinate (probability) and y-coordinate (magnitude) of a two-dimensional plot (Figure 1E, F, G). The IPs of all equally frequently chosen options align as an indifference curve (IC) irrespective of their different magnitude-probability composition. The colored curves in Figure 1F and G are ICs. As we inferred subjective economic value from observable choice, we assumed that all options on the same IC have the same subjective economic value, and options on different ICs have different value. Options on higher ICs (farther from origin) are more frequently chosen than options on lower ICs, and thus have higher value. For example, options on the cyan and pink ICs in Figure 1F, G are more frequently chosen, and have higher value, compared to options on the amber and green ICs.

All data collection and analysis were performed using custom code in MATLAB (version 8.3.0 (R2014a). Natick, Massachusetts: The MathWorks Inc).

### Using the continuity axiom for value estimation

Expected Utility Theory proposes four axioms that determine the maximization of utility (von Neumann and Morgenstern, 1944). Compliance with completeness (axiom I) and transitivity (axiom II) is necessary for consistently ranking all choice options. Compliance with continuity (axiom III) demonstrates that choices reflect a meaningful representation of numerical utility. The independence axiom (axiom IV) defines the computation of expected utility from reward magnitudes and their probabilities. The four axioms of EUT define necessary and sufficient conditions for choices to be described by the maximization of subjective economic value: if the axioms are fulfilled, a subjective economic value can be assigned to each choice option, and the decision maker behaves as if choosing the highest valued option (von Neumann and Morgenstern, 1944). The third axiom, continuity, requires the testing of a wide range of reward magnitudes and probabilities. The extended testing in conceptually well-defined conditions seemed particularly helpful for detecting unreliable neuronal coding. Therefore, the continuity axiom with its extended tests constitutes the foundation of the current study. The continuity axiom is stated as:

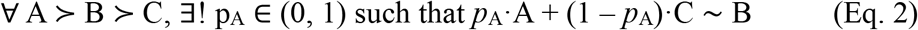

where A, B, C and AC are gambles, “≻“ is the preference relation, and “~” the indifference relation. In words, compliance with the continuity axiom requires to identify a reward probability *p*_A_ at which choice indifference occurs between a fixed gamble B and a variable gamble AC. The variable gamble AC consists of a probabilistic combination of a higher valued gamble A (with probability *p*_A_) and a lower valued gamble C (with probability 1 - *p*_A_) (Eq. 1). Compliance with the axiom implies the possibility of defining a numerical scale of subjective economic value. As a deterministic rule, the axiom assumes constant preferences over time. As neuronal statistics requires repeated choices and as neuronal responses are inherently variable, we interpreted the axiom in a stochastic sense: option A was considered stochastically preferred to option B when the probability of choosing A over B was larger than 0.5 (binomial test, *p* < 0.05).

According to the continuity axiom, the behavioral indifference point (IPb) represents a utility measure, being a numerical quantity associated with the subjective evaluation of gamble B in relation to gambles A and C. Following the formalism of the continuity axiom, we defined subjective economic value behaviorally as the estimated reward probability for which the monkey was indifferent between the safe option B and the combined gamble AC (Figure 1C). The IPb was computed according to a standard discrete choice model; we fitted a softmax function to the probability of choosing the AC option and identified the point for which the softmax’ value was 0.5, which corresponded to a 0.5 probability of choosing equally frequently each option, i.e., choice indifference.

The numerical value of the indifference point (IP) in choices between the safe option B and the combined gamble AC was determined by fitting a softmax function to the choice data through non-linear least squares fit:

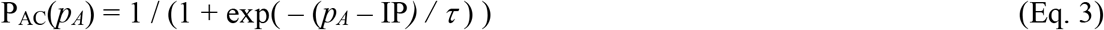

with IP corresponding to the probability *p_A_* resulting in equal preference for the two options, and *τ* as softmax ‘temperature’ parameter representing the steepness of the choice function (steeper for lower *τ* values).

### Behavioral economic model fit for subjective economic value

The subjective economic value of a gamble derived from reward magnitude *m* and probability *p* and was captured by the product of a utility function *u(m)* and a probability weighting function *w(p)*:

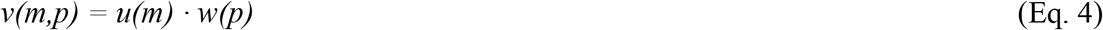

with *m* as magnitude (in ml), *p* as probability (*p* ∈ [0,1]), and *u* and *w* as two separate 2-parameter *Prelec* functions (Prelec, 1998):

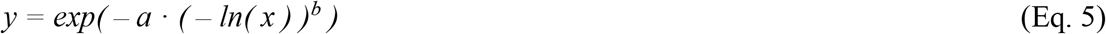

with *y* as generic Prelec function that gave rise to the 2-parameter utility function *u(m)* and the 2-parameter probability weighting function *w(p).* The Prelec function’s *x* was defined between 0 and 1, while magnitude levels varied between 0 and 0.5 ml; therefore, in the *u(m)* we set *x* = *m* / 0.5.

We modeled the probability of choosing one option using a standard discrete choice model. The probability of choosing gamble A in choices between any two gambles (option set {A,B}) was defined through a binary logistic model:

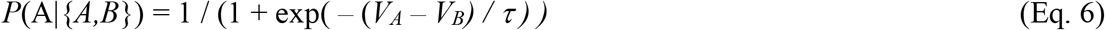

with *V_A_* and *V_B_* as the subjective economic values of gamble A and B, respectively, and*τ* as the softmax temperature. The gamble values V_A_ and V_B_ were defined according to Eq. 4.

The two parameters (*a* and *b*) of the Prelec utility function *u(m),* the 2 parameters of the Prelec probability weighting function *w(p)*, and the softmax temperature*τ* were estimated by maximum likelihood estimation (MLE). The MLE procedure involved computing and maximizing the log-likelihood (*LL*) in the parameter space:

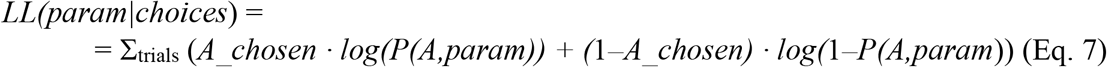

with the sum Σ being defined across all trials in one session, *A_chosen* as the value of 1 if gamble A was chosen in one trial and 0 otherwise, and *P(A,param)* as the discrete choice model defined above with parameters *param.* We minimized the negative LL using the *fminsearch* Matlab function.

To obtain an IC for a given fixed option B, we estimated a series of equally frequently chosen IPb’s. Each IPb was obtained by varying the reward probability in gamble A. Each IC was obtained by setting a series of reward magnitudes for gamble A. Thus, we obtained a different IC for each magnitude of gamble B. As by definition all equally frequent choice options were located on the same IC, we inferred that all IPb’s on a given IC had the same subjective economic value. Points on an IC for a higher-magnitude B had higher subjective economic value.

We derived an analytical expression for the ICs in order to represent any IC in the magnitude-probability graph using the four fitted parameters. As by definition all IC points have the same economic value, the general equation for an IC was obtained by equating the economic value of a fixed safe option B (with magnitude m_B_) with the economic value of a generic gamble G (with magnitude m and probability p). Because V_B_ = u(m_B_) and V_G_ = u(m) · w(p), the equation corresponds to

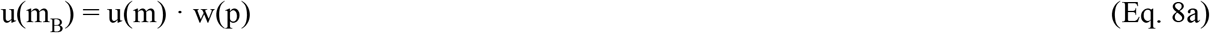

from which we extracted p(m), i.e., an expression for all points (m, p) with economic value equal to the value of a safe option B.

Formally, we introduced the parametric expression for the utility (u) and probability weighting (w) functions (Eq. 5)

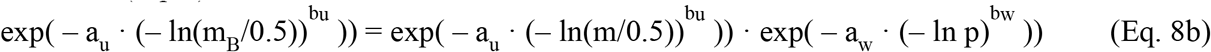

isolated the p term after applying the logarithm to both sides

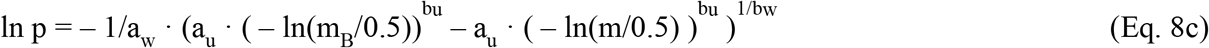

and derived the IC equation by applying the exponential to both terms:

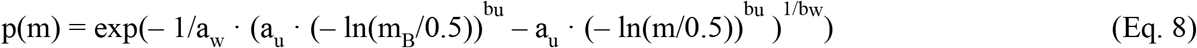

This curve represents the relation between probability and magnitudes along the indifference curve passing through the point (1, m_B_), when utility and probability functions are defined as Prelec functions.

### Neurophysiological recordings

A head-restraining device and a recording chamber were implanted on the skull under full general anesthesia and aseptic conditions. The chambers were 35 mm wide by 40 mm long (Monkey A) and 40 mm wide by 50 mm long (Monkey B; Gray Matter). The stereotactic coordinates of the chamber enabled neuronal recordings of the orbitofrontal cortex (OFC). We located the OFC from bone marks on coronal and sagittal radiographs taken with a guide cannula inserted at a known coordinate in reference to the implanted chamber, aiming vertically for areas 11 and 13 (central orbital gyrus; stereotaxic coordinates: 27 - 37 mm anterior to interaural line, 3 - 12 mm lateral to midline) (Figure S2). Monkey A provided data from the right-hand hemisphere and Monkey B from both hemispheres via a craniotomy ranging from Anterior 28 to Anterior 38 and Lateral from 2 to 12. We conducted single-neuron electrophysiological recordings using glass-coated tungsten electrodes, custom made or commercial (Alpha Omega), with impedance of about 1 MOhm at 1 kHz. One or two microelectrodes were inserted into the cortex with a multi-electrode drive (NaN instruments). Neuronal signals were collected at 20 kHz, amplified using conventional differential amplifiers (CED 1902 Cambridge Electronics Design) and band-pass filtered (high: 300 Hz, low: 5 kHz). We used a Schmitt-trigger to digitize the analog neuronal signal online into a computer-compatible TTL signal. However, we did not use the Schmitt-trigger to separate simultaneous recordings from multiple neurons, in which case we searched for another recording from only a single neuron, or we stored occasionally the data in analog form for off-line separation by dedicated custom-written software in MATLAB. An infrared eye tracking system monitored eye position (ETL200; ISCAN), with temperature check on an experimenter’s hand at the approximate position of the animal’s head.

### Neuronal task relationships

The main data of this study were obtained in a zero-alternative task in which the animal was presented with either the safe option B or the gamble AC. When a single neuron was identified and isolated, the zero-alternative task contained a limited set of reward parameters (safe option B: 0.15 ml or 0.35 ml, probability 1.0; gamble AC: magnitude 0.5 ml, probability 0.3 or 0.7). Each option was presented 8 times, resulting in a total of 24 trials. We selected OFC neurons according to their significant response to the Cue (P < 0.05; Wilcoxon test comparing neuronal activity during a 400 ms time-window starting at Cue onset vs. baseline activity during 400 ms immediately preceding the Cue). Then we ran the full set of reward parameters on these significantly modulated neurons. These parameters resulted in 7 – 20 zero-alternative trial classes (at least 3 safe options and 4 gambles) intermingled and equally presented for at least 8 repetitions (12 replications on average). In a subset of sessions, we also presented 6 – 18 binary-choice additional trial classes, all intermingled with the zero-alternative ones. Additional binary-choice trials were collected between recording sessions and while moving the electrodes in search for isolated neurons (for the collection of behavioral data on the same day of the neuronal recordings in as many tests as possible). We interrupted the session when a neuron’s waveform became too small or was lost during the recording. We included neurons that were well-isolated for at least 4 repetitions. The single-neuron activity in one trial was calculated as the rate of action potentials after the Cue, expressed as neuronal impulses per second (imp/s). When offline-sorting the signals, other neurons not observed during recordings were added to the database if their waveform was clearly different from that of other simultaneously isolated neurons and well separated from the background noise signal.

In order to identify basic reward magnitude and probability coding, we selected OFC neurons for their Cue responses varying with both reward magnitude and reward probability, as defined by significance of the multiple linear regression:

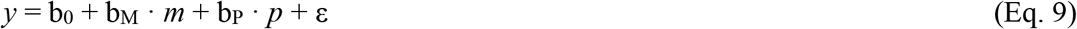

with *y* as neuronal activity (imp/s) during the 400 ms time-window after the Cue, b0 as offset, bM and bP as slopes for the regressors of magnitude (*m,* in ml) and probability (*p* ∈ [0,1]), respectively, ε as error term. In summary, for this initial analysis, we considered only OFC neurons whose Cue responses varied significantly against baseline (P < 0.05; Wilcoxon) and encoded both reward magnitude and reward probability as demonstrated by both slope parameters b_M_ and b_P_ being significantly different from 0 (P < 0.05; t-test).

The analysis time-window after the Cue included the initial part of the immediately following choice movement and thus did not provide sufficient temporal difference for separate analysis; we did not analyze responses to the reward due to its lack of explicit probability information.

For comparison with the data obtained in the zero-alternative task, we recorded OFC neurons during binary choice. Such neurons fall broady into two classes of interest for the current study; chosen value neurons code the value of the particular option the animal chooses (value coding depending on the animal’s choice); object value neurons code the value of only one choice option regardless of the animal’s choice (object value is also called offer value; Padoa-Schioppa and Assad, 2006; a third neuron type identified by these authors refers to the choice in a binary manner, rather than to value, and was therefore not considered for our study).

The two framed options in our task were visually distinguishable and allowed unambiguous identification of each option as requirement for distinguishing between chosen value coding and object value coding. Option AC (probabilistic gamble) was indicated by 2 graded horizontal value bars whose lengths provided explicit information about the probability of each reward magnitude, whereas option B (‘safe’ reward) was indicated by a single bar extending over the full width of the quantitative stimulus (Figures 1A, B; 2B).

We identified chosen value neurons using the following regression:

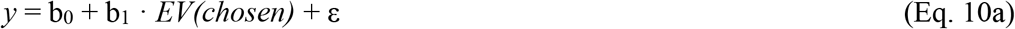

to be classified as chosen value coding, the slope parameter b_1_ needed to be significant for the chosen option’s EV (P < 0.05; t-test). We excluded neurons coding the value of the unchosen option, which in our design correlated with the value of the chosen option; we also excluded neurons coding the value difference between the two choice options. We identified neuronal responses coding the value of the unchosen option using the following regression:

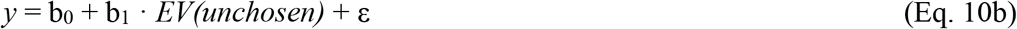

and we identified neuronal responses coding the value difference using the following regression:

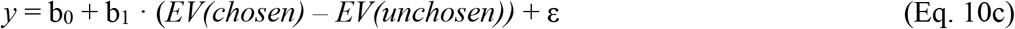

We only included as chosen-value neurons those with a higher r-squared for the chosen-value regression compared to the r-squared of both the unchosen-value regression (Eq. 10b) and the value-difference regression (Eq. 10c).

We identified object-value coding neurons as those coding only the value of one of the two choice options (gamble AC or safe reward B) regardless of the actual choice the animal made. This was assessed using two separate linear regressions of the neuronal activity with either the expected value (EV) of the gamble option, or the EV of the safe option (P < 0.05). We identified object value neurons using the following regressions for the gamble and safe options (significant slope coefficient b_1_):

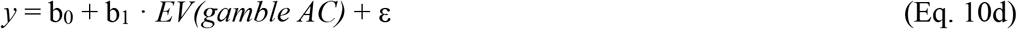

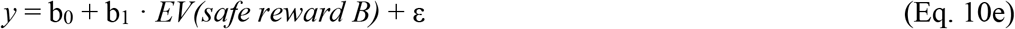

### Neuronal subjective economic value measure

During neuronal recording sessions we presented choices between one option (gamble or safe) and no reward. These zero-alternative trials were intermixed with binary choices between a gamble and a safe option or presented in separate sessions. The zero-alternative task represented a fully controlled situation for studying the neuronal responses to one specific option, and thus to investigate the neuronal coding of value without confounds introduced by varying, simultaneously presented alternative options. We present neuronal data from the zero-alternative trial types, in which information was only relative to the non-zero option. We then use the same analysis for gamble-safe choice trials, in which we characterized the neuronal activity as a function of the reward variables (*m* and *p*) relative to the chosen option.

### Elicitation of neuronal indifference points

Mirroring the behavioral utility measure, we defined a neuronal subjective economic value measure for a particular set of reward magnitudes {A, B, C}. The neuronal indifference point (IPn) corresponded to the probability of A for which the neuron had the same response to the gamble (AC) and to the safe option (B). The IPn was computed by first decoding the probability of choosing the gamble AC from the neuronal activity (see below). Then, a softmax function was fitted to the decoded choice probability, identifying the point at which the softmax had a value of 0.5 (i.e., decoded choice indifference) (Figure S4).

### Neuronal noisy representation of behavior

To test whether the neuronal activity represented the behavior, we performed two data analyses. First we looked for neurons with significantly different activity (t-test, P < 0.05) in response to two equally preferred options. Second, we investigated whether the recorded activity was compatible with the noisy coding of behavior, assuming Poisson-distributed activity rates.

In the first analysis, we selected all neurons recorded in the same four tests. We then compared the activity in response to the two options (safe and gamble) that were closest to the ones resulting in behavioral indifference. To compute the percentage of neurons deviating from behavior we randomly selected (20,000 iterations) one of the tests and counted the neurons with significantly different activity (t-test, P < 0.05). We repeated the analysis with random selection of two, three or four tests, counting the number of neurons with different activity in at least one test. Neurons were also randomly selected with replacement, therefore the error bars (Figure 4A) represented variations both across tests and across neurons.

The second analysis investigated whether the recorded activity was compatible with the noisy coding of behavior. As the cortical activity is commonly assumed to be Poisson-distributed, we simulated the activity of neurons coding the economic value assuming Poisson-distributed activity rates. The response to each tested option was simulated as a noisy Poisson process with mean rate proportional to the economic value of that option. The simulation was repeated for each neuron; the baseline and range of simulated activities was set equal to the mean ones of the corresponding neuron. The simulation was implemented using a standard computational algorithm (Dayan and Abbott, 2005). We focused on three option pairs (safe, gamble) with equal EV, away from behavioral indifference, corresponding to EV of 0.15 ml, 0.25 ml and 0.35 ml respectively. To test if a neuron’s activity was compatible with the simulated one, we computed the activity difference between the two options and the 95% confidence interval of the same difference for the corresponding simulated neuron (2,000 iterations). We then counted the neurons whose mean activity difference fell outside of the simulated 95% range.

### Choice-probability decoder

To directly relate the neuronal activity to a behavioral quantity, we decoded preferences from the neuronal activity and compared them with the measured behavioral ones. To decode preferences, defined as the percentage of choices for one option over another option, we used a specifically designed choice-probability decoder, closely related to the Receiver Operating Characteristic analysis of signal detection theory (Green and Swets, 1966; Britten et al., 1996). Assuming that the activity of a neuron was proportional to the instantaneous subjective value of an option, including any noise in the system, a higher activity would identify the preferred option. Across multiple trials, the strength of preference would then be related to the overlap between the distributions of activity rates for the two options (Figure S4A, B). Higher preference (indicated by ≻) was defined as the probability of choosing one option (X) over the other option (Y) (P(choice) > 0.5). Higher neuronal preference was defined in analogy as percentage of trials with higher activity for option X than for option Y. Thus, the probability of choosing option X over option Y coincided with the percentage of trials with higher activity for option X than for option Y (see Figure 4C, D):

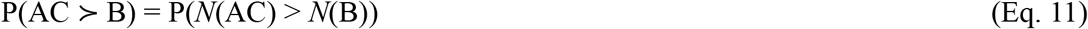

where *N*(G) represents the distribution of activity rates of a neuron when presented with gamble G.

Based on this definition, we computed the probability of choosing option AC as the probability that the activity for AC was higher than the activity for B. We counted the number of occurrences in which the activity for AC was higher than the activity for B, across all possible pairings of single AC and B trials (which were collected independently, in the zero-alternative task). The decoded choice probability was then calculated as the number of occurrences divided by the total number of pairings.

This method allowed us to represent the neuronal and behavioral quantities on the same scale (P(choose AC) ∈ [0,1]), for a direct comparison of the behavioral and neuronal indifference point measures (Figure 2D).

Moreover, we used the neuronal preference measure to characterize neurons in relation to behavior in terms of their global risk attitude coding (Figure 4C). The decoded neuronal preferences (P_N_) were regressed on the behavioral preferences P_B_:

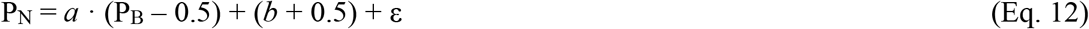

where the regression slope (*a*) quantified the strength of preference coding, while the intercept (*b* + 0.5) represented the relation with the behavior. Neurons for which *b* = 0 would consistently encode the behavioral risk attitude across tested gambles; neurons for which *b* < 0 (or *b* > 0) would encode a more risk averse attitude (or a more risk seeking one) than behavior.

In this regression, although behavioral preferences act as the regressor and neuronal preferences as the response variable, we did not imply a relation of dependence between neuronal and behavioral data. The significance of the slope parameter is used as a signature of the correlation between the variables (the P value for the slope parameter is equivalent to that of a correlation coefficient), and the intercept is needed to define different encoded risk attitudes.

### Neuronal economic model fit for subjective economic value

We fitted an economic multiplicative value model directly to the neuronal activity recorded in all trials. The multiplicative model represented the subjective economic value of a gamble as the product of a utility function and a probability weighting function, as in the behavioral model, using a 2-parameter Prelec function (Eqs. 4 and 5). To do so, we fitted the 2-dimensional value function to the neuronal activity *N* through nonlinear least squares:

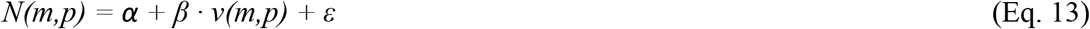

with free parameters *α* (baseline activity), *β* (activity range) and *ε* (error term), using data from all presented gambles. The constraints imposed to the procedure (using Matlab function *fminsearchbnd*) imposed *N* > 0, ensuring that meaningful neuronal activities were produced by the model. Additional constraints were imposed for biological plausibility and fit stability by limiting the modeled activity to not exceed the recorded activity range by 30% (this controlled for implausibly large values). This analysis was applied separately to each neuron.

### Neuronal population code

We defined a population-estimate of decoded preferences following the heuristic argument that, in noisy multi-component systems, accuracy can be improved by computing the output signal based on the majority of outputs from single components (von Neumann, 1956). Preference was defined as the proportion of gamble choices in a gamble-safe choice pair (PchAC). Assuming that multiple neurons encoded the same preference with independent noise contributions, the choice encoded by the majority of neurons would drive the population-level preference in that direction. Thus, we computed the resulting preference value as the percentage of neurons encoding a preference for the gamble option (i.e., corresponding to PchAC > 0.5).

To compute the coding of economic functions by a neuronal population, we computed the median value of the estimated economic functions’ parameters (2 parameters for the Prelec function for utility, *u(m),* 2 parameters for the Prelec function for probability weighting, *w(p)).* We computed the economic functions from a neuronal population by computing the median of each parameter across the neurons. The median was used to include the effect of asymmetric parameters’ distributions, and to reduce the dependency on outliers.

The procedure was iterated in a bootstrap procedure (resampling with replacement; n = 20,000 iterations). At each iteration, the utility function and probability weighting functions were computed from the median parameters, and then averaged to obtain the populations’ functions. The 95% confidence intervals for the neuronal economic functions were defined by selecting the 2.5^th^ to the 97.5^th^ percentile for each x-value) (Figure 6B). The same procedure was applied to each IC, separately (Figure 6C).

## QUANTIFICATION AND STATISTICAL ANALYSIS

All statistical tests (t-test, Wilcoxon test, correlation (Pearson), binomial test) were considered significant at the α = 0.05 level.

We used a bootstrap procedure to compute the error areas associated with utility and probability weighting functions in single neurons (Figures 5B, S7B, S9A). We resampled with replacement the activity of single trials and then fitted the economic value functions to the resampled data. The procedure was repeated 2,000 times and the standard deviations and 95% confidence intervals from the resulting curves were computed. The same procedure produced the error area around each IC (Figure 5C; Figure S7C, S9B).

To test if the measured percentage of neurons yielding IPn close to IPb was significantly different from the prediction of a random model (i.e., a model assuming that IPn’s were noisy representations of the IPb’s), we used a Monte Carlo simulation (Buckland, 1984). We assumed normal distributions for the IPn values equal to the ones estimated from the IPb values. After the simulation was run (n = 10,000 iterations), the results were computed by calculating the simulation percentages and 95% confidence intervals (from the 2.5^th^ to the 97.5^th^ percentile) across iterations (Figure 4A).

## Supplemental Information

**Figure S1.**
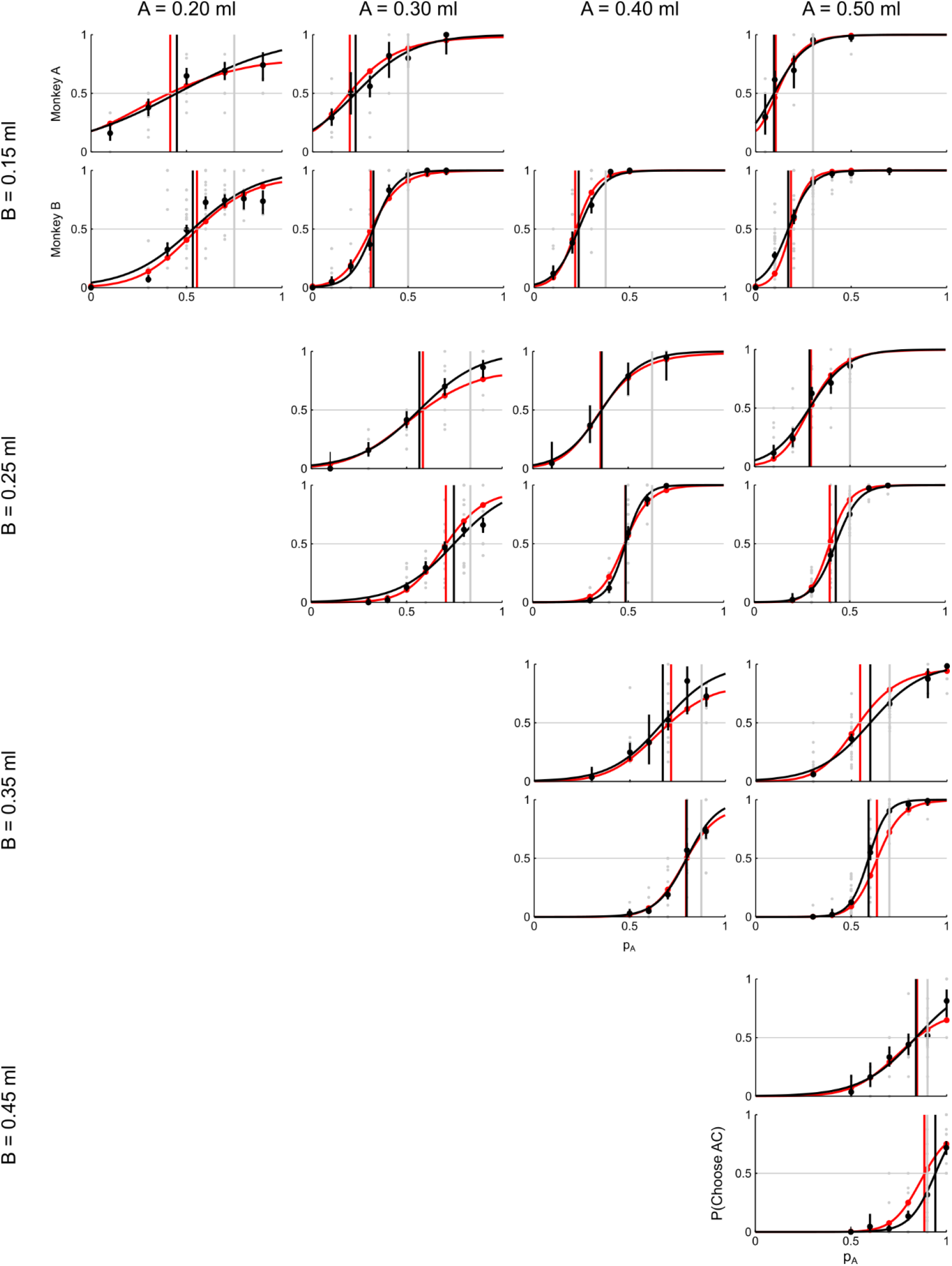
Behavioral data and economic model. Comparison between behavioral data (black, across all sessions) and economic model prediction (red). The fitted model (Eq. 6) was able to capture the indifference points (vertical lines) as well as the predicted preferences (P(choose AC), y-axis) across the wide range of tested magnitude and probability levels. Gray: single sessions’ preferences.

**Figure S2.**
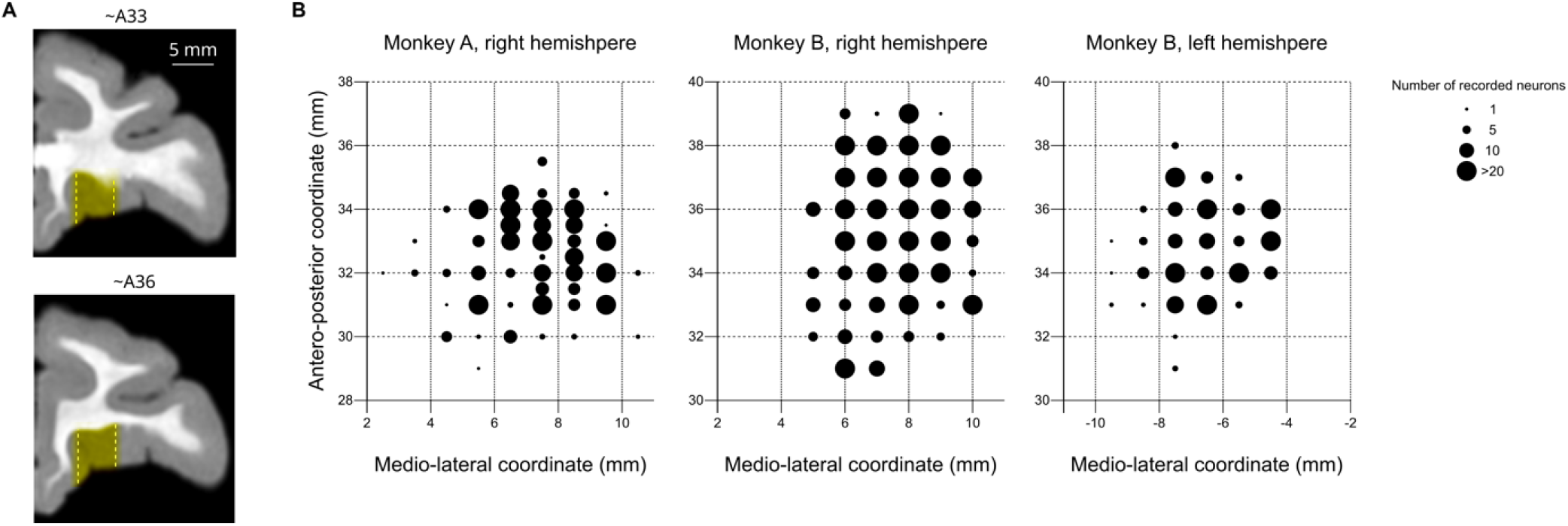
Neuronal recording locations. (A) Recording regions. The majority of recording locations were between 5 and 10 mm lateral coordinate (dashed lines), implying that the recorded neurons were mostly located in the central/medial orbitofrontal gyrus. (B) Numbers of recorded neurons per location, in three hemispheres of both monkeys. Anteroposterior coordinates are relative to the ear bars.

**Figure S3.**
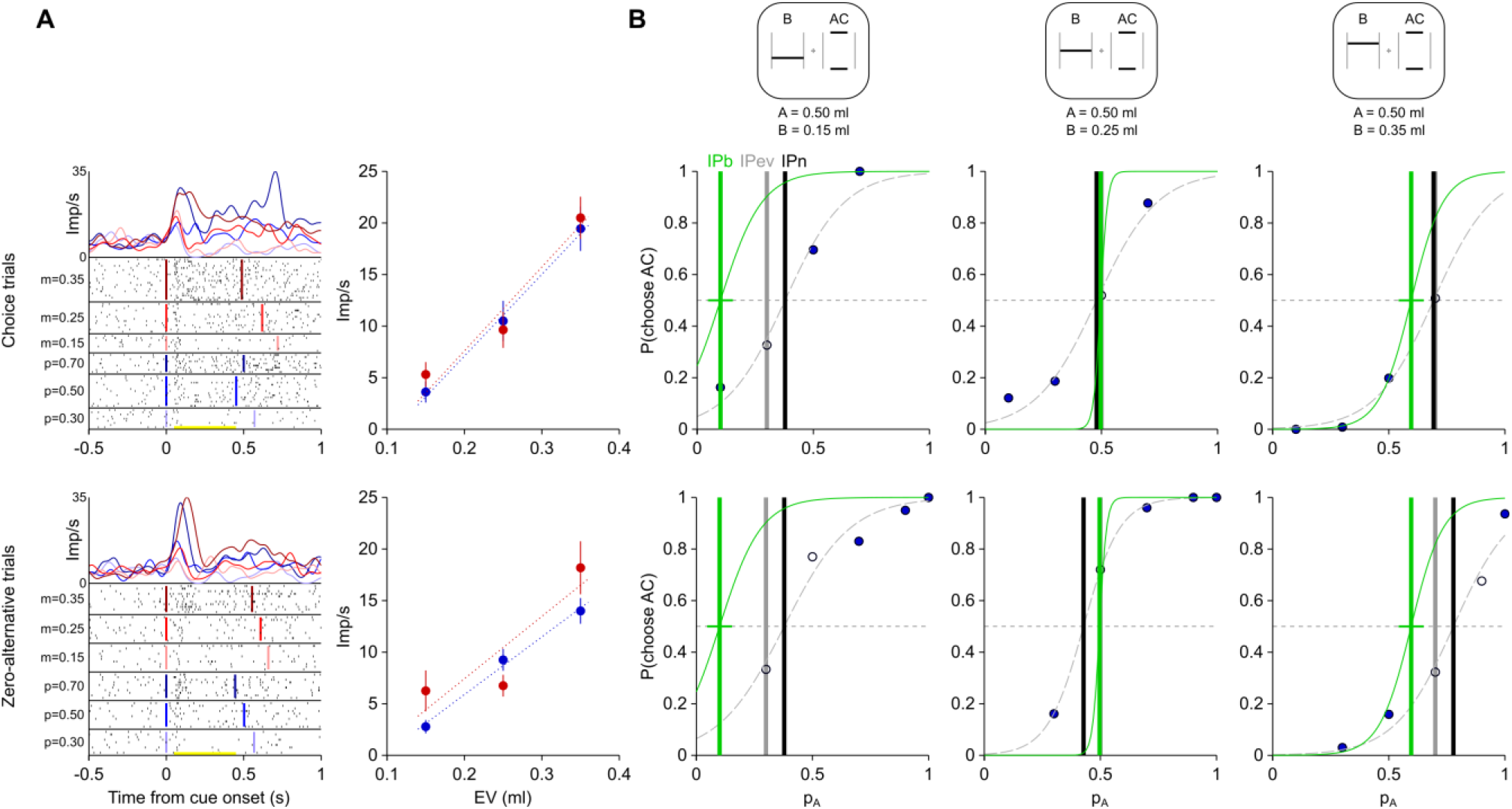
Chosen value coding in single OFC neuron. Reward magnitude and probability coding in a single OFC neuron in choice trials. Top: choice trials, bottom: zero-alternative trials for comparison; Monkey B. For conventions, see Figure 2. (A) Left: response to the chosen option indicated on the y-axis (*m:* magnitude of safe option B, *p:* probability of gamble AC). Right: average (± SEM) activity rates computed in the analysis window (yellow bar in the raster plot). Linear fits (dotted lines) show sensitivity of neurons to both reward magnitude and probability. Imp/s: impulses per second. (B) Corresponding variations of neuronal indifference points (IPn, black vertical lines) between choice and zero-alternative trials. IPb: behavioral indifference point (green vertical line), IPev: p_A_ for which EV(B) = EV(AC) (p_A_: probability of gamble A in the gamble mix of A and C).

**Figure S4.**
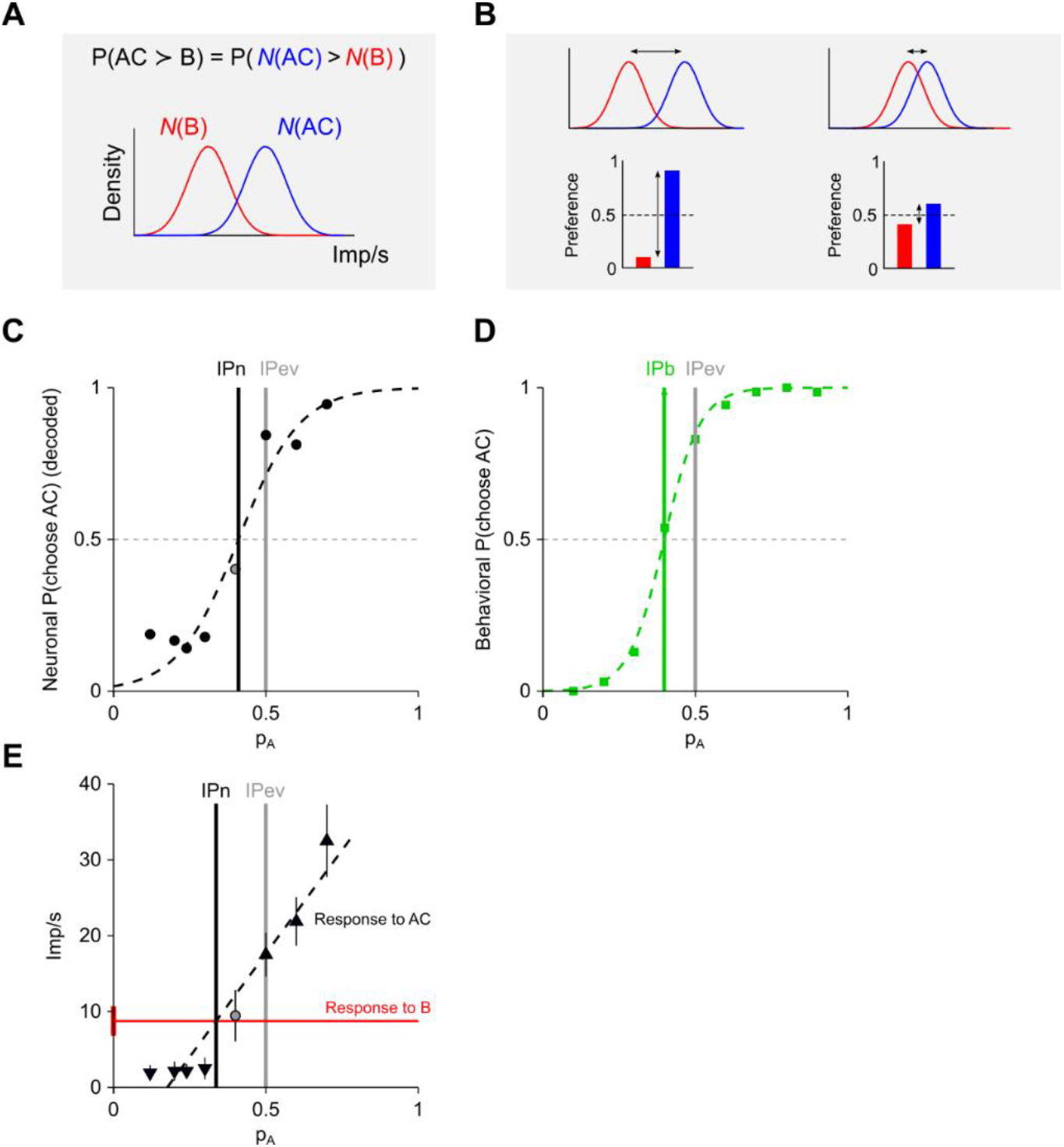
Preferences decoded from the activity of an individual neuron. (A) Scheme of the choice-probability decoder. For a value coding neuron, the trial-by-trial activity would be proportional to the instantaneous subjective value of an option, including any noise in the system. A higher activity would thus identify the preferred option. Across multiple trials, the strength of preference would then be related to the overlap between the distributions of activity rates for the two options. The probability of choosing option AC over option B (i.e., the probability that option AC had a higher value than option B) coincides with the percentage of trials with higher activity for option AC than for option B. (B) Example decoded preferences. The overlap between the distributions of neuronal activity for trials presenting option AC and option B determine the decoded preferences. Preferences are high (P ~ 1) when the two distributions are well separated (left), while they are close to indifference (P ~ 0.5) when the two distributions are mostly overlapped. (C), (D) The neuronal decoded preferences (C) can be used to infer the neuronal subjective value measure (IPn) of the neuron, and can be represented in the same graph as the behavioral preferences, for direct comparison (D). (E) A simpler method for computing of the IPn values, without decoding preferences, involves the direct comparison of activity rates. The IPn can be defined as the probability p_A_ for which a linear regression of the activity for options ACs (dashed line) intercepts the mean activity for option B (red line). This method gives similar results in terms of heterogeneity between OFC neurons (not reported) but involves some crucial limitations: it assumes linear responses to reward probability, it relies on the computation of a single value for the activity of B, and results cannot be represented in the scale of the behavioral preferences (*p* ∈ [0 1]). For these reasons, we did not use this method for our data analysis. Imp/s: impulses per second.

**Figure S5.**
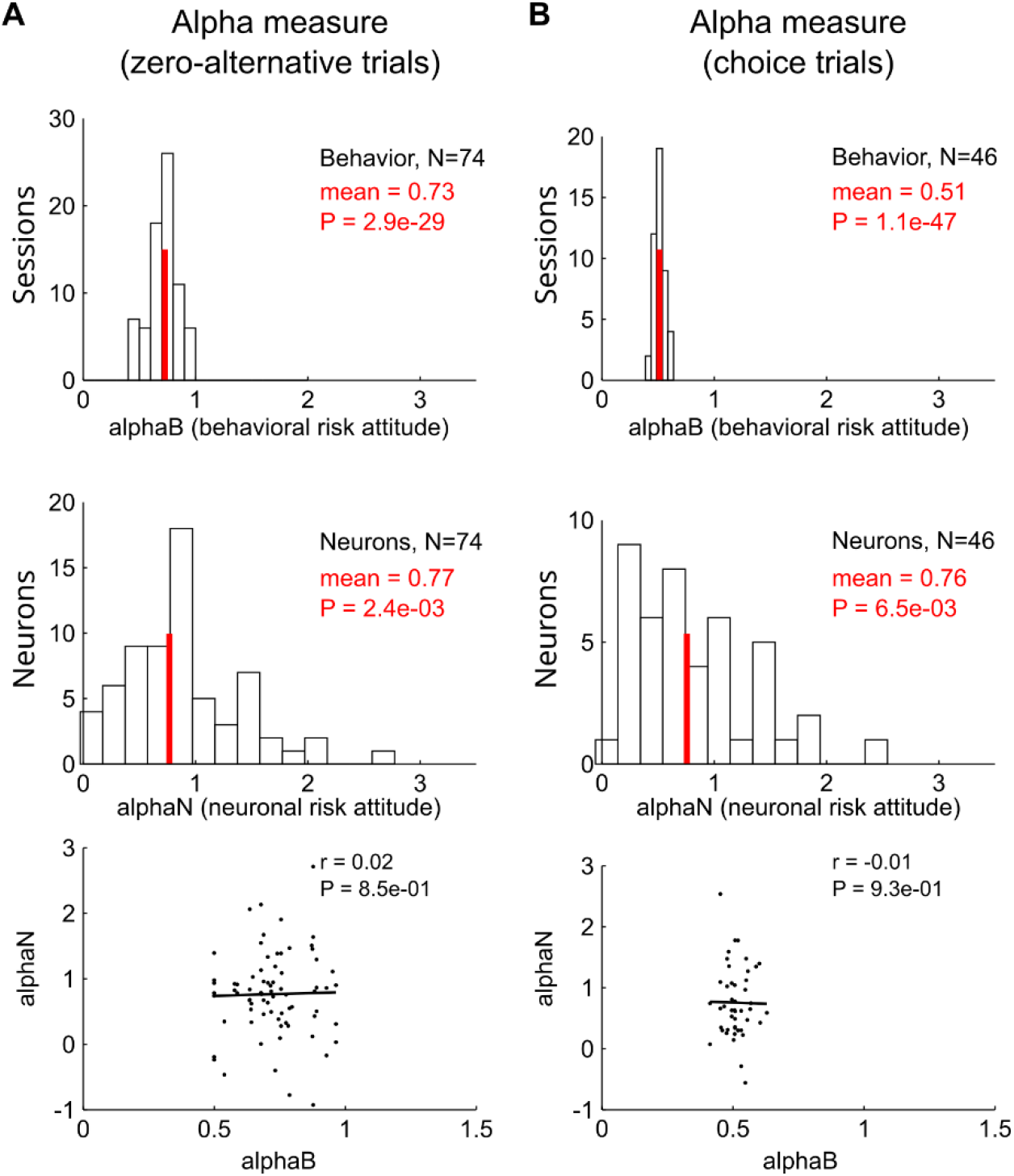
Correlation between neuronal and behavioural risk-attitude measures. (A) Risk attitude measure (alpha) as defined in Raghuraman & Padoa-Schioppa 2014. Top: histogram of behavioral measure values (alphaB). Center: neuronal measure (alphaN). Bottom: correlation between neuronal and behavioral measures (r: Pearson’s correlation coefficient, with p-value P). Data from Monkey B during zero-alternative trials. Red line: mean value (P: p-value of *t-* test against 1). (B) Alpha measure during choices. Data from Monkey A.

**Figure S6.**
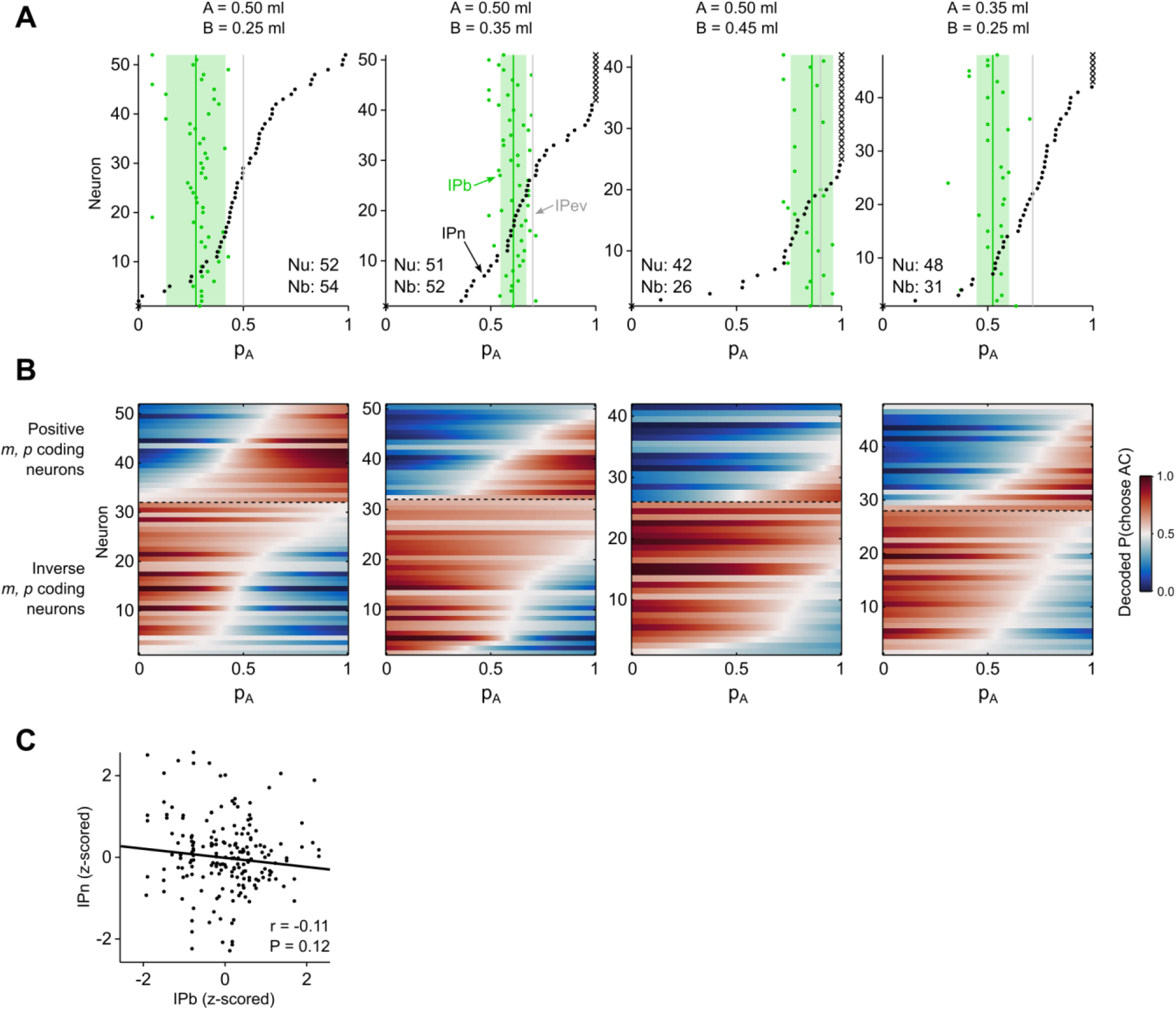
Neuronal subjective value measure during choices. Subjective value representation in a binary-choice task. Conventions and symbols as in Figure 3. Neuronal activity was analyzed in relation to the magnitude and probability of the chosen option. Neurons were recorded during choices between options spanning at least 2 safe magnitudes, 3 gamble probabilities and 2 gamble magnitudes.

**Figure S7.**
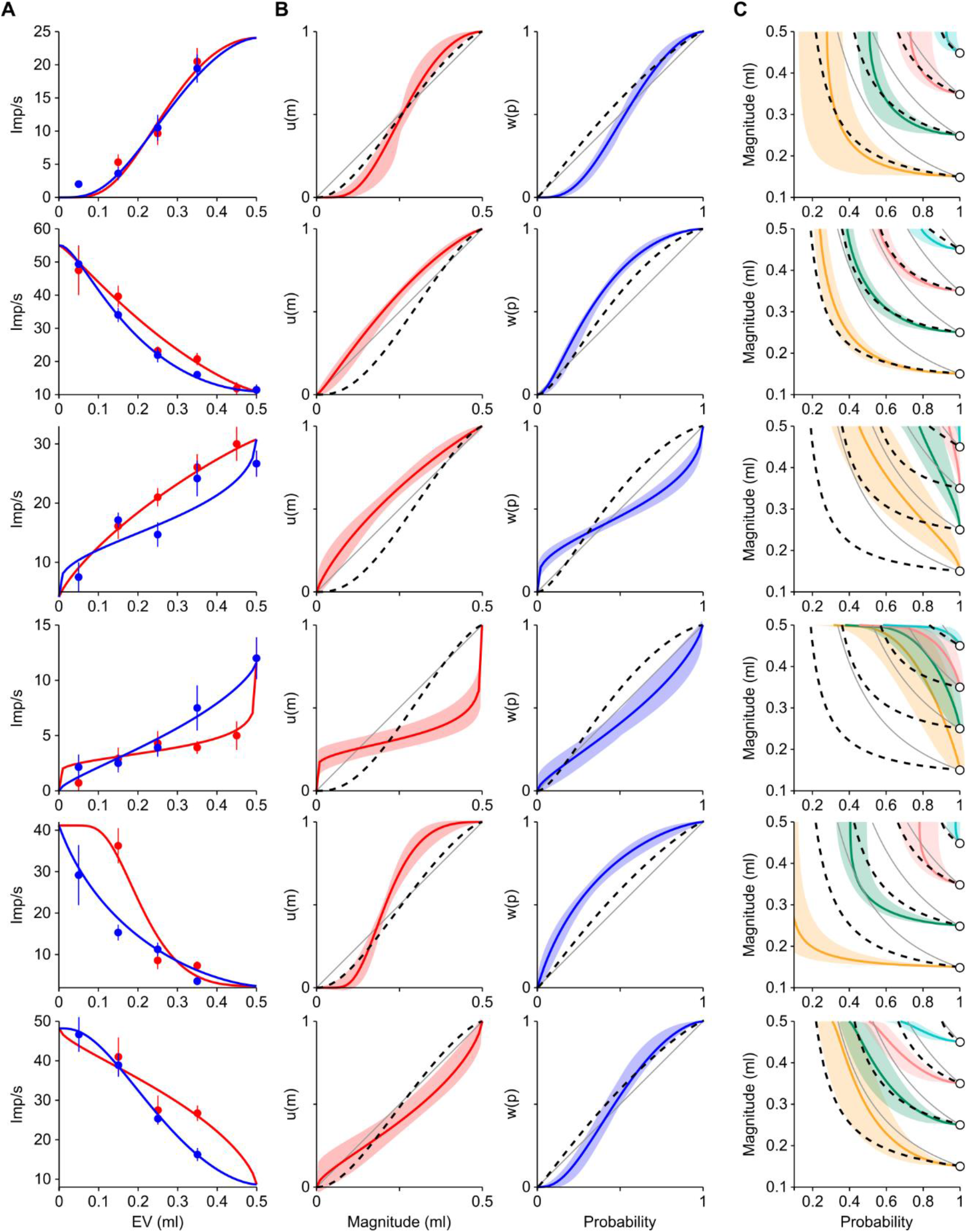
Diverse coding of economic value in chosen value neurons. Six example neurons highlighting the heterogeneous coding of economic value during choice, in terms of activity pattern (panel A), reconstructed utility and probability weighting functions (panel B), and indifference curves (panel C). Each row shows data from the same neuron (rows 2, 3, 4: neurons from Monkey A; rows 1, 5, 6: neurons from Monkey B). Conventions as in Figure 5A-C.

**Figure S8.**
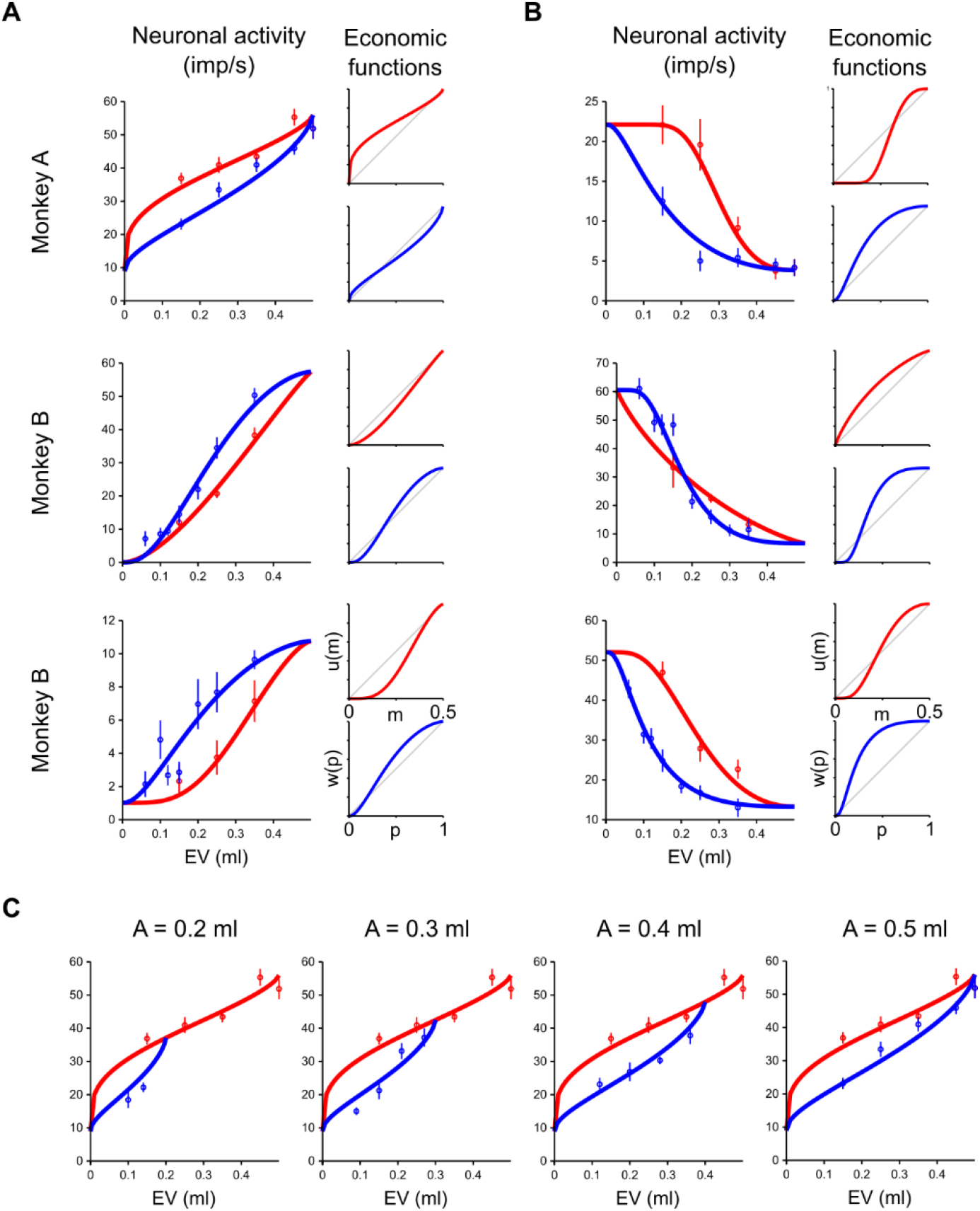
Single neurons’ example responses and economic functions. (A) Example positive *m, p* coding neurons. Left: response to different reward magnitudes (red) and probabilities (blue) (dots: mean; vertical bars: SEM), revealing different nonlinear response patterns across neurons, in both monkeys. The curves were obtained by fitting the subjective economic value function to the neuronal data (Eq. 13). Right: best-fitting utility function (red, top) and probability weighting function (blue, bottom). Imp/s: impulses per second. (B) Example inverse *m, p* coding neurons (same data representation as in panel A). (C) Activity and fitting value functions for different magnitude levels for gamble A. A single set of fitting parameters explained the activity for multiple gamble magnitudes and probabilities (the economic model fit to the neuronal activity included all tested *m* and *p* levels).

**Figure S9.**
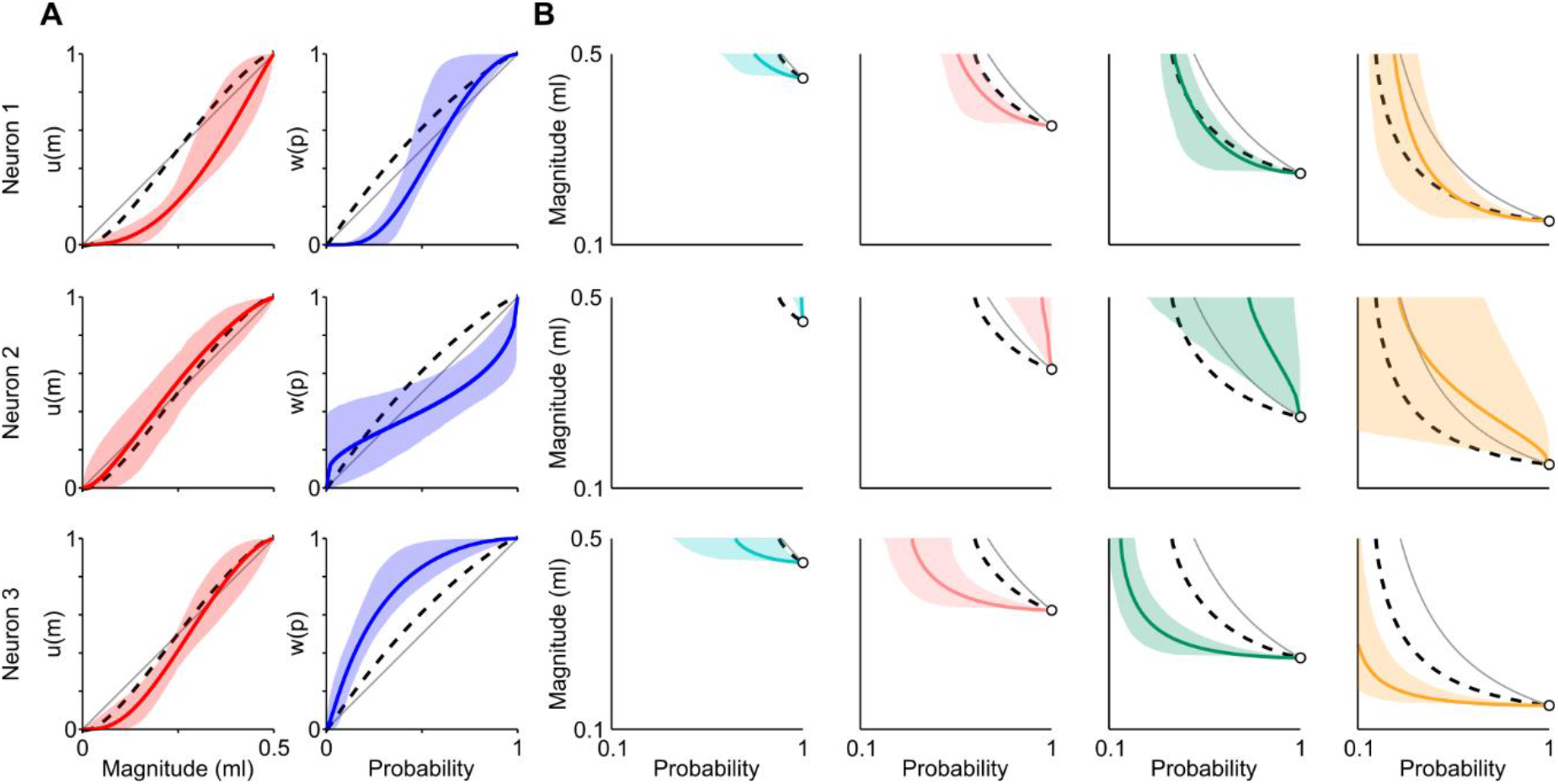
Statistical comparison between neuronal and behavioral data in single neurons. (A) Utility (left) and probability weighting (right) functions recovered from behavior (dashed lines) and from the activity of individual neurons (colored lines), with 95% confidence interval (bootstrap). (B) Indifference curves recovered from behavior (dashed lines) and from the neuronal activity (colored lines), with 95% confidence interval (bootstrap). Grey lines: EV-curves. Conventions as in Figure 5.

**Figure S10.**
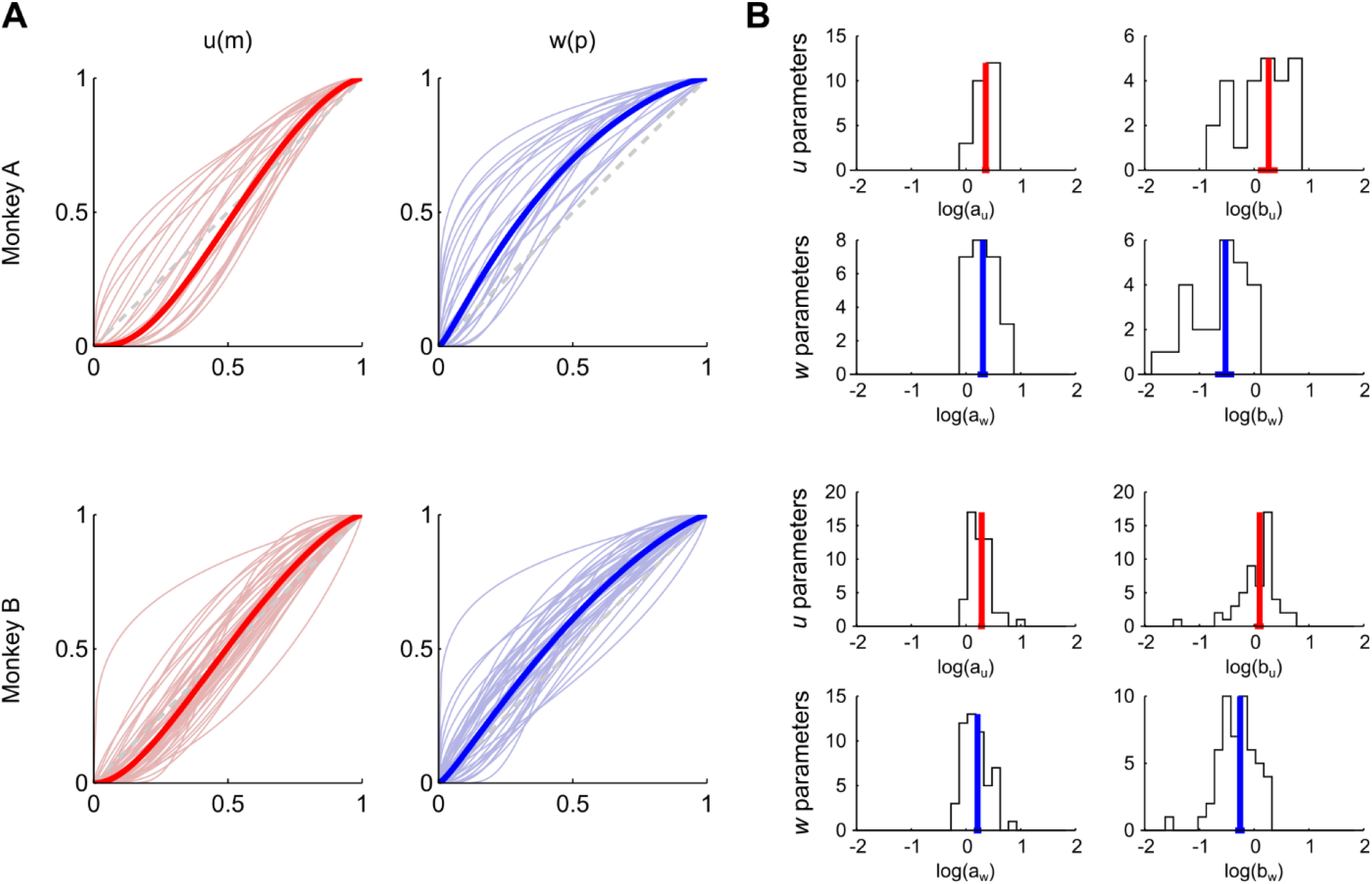
Utility and probability weighting functions in single behavioral sessions. (A) Utility function (left, red) and probability weighting function (right, blue) for single behavioral sessions (thin lines) and corresponding mean functions (thick lines). (B) Distribution of *u* and *w* parameters across behavioral sessions.

**Figure S11:**
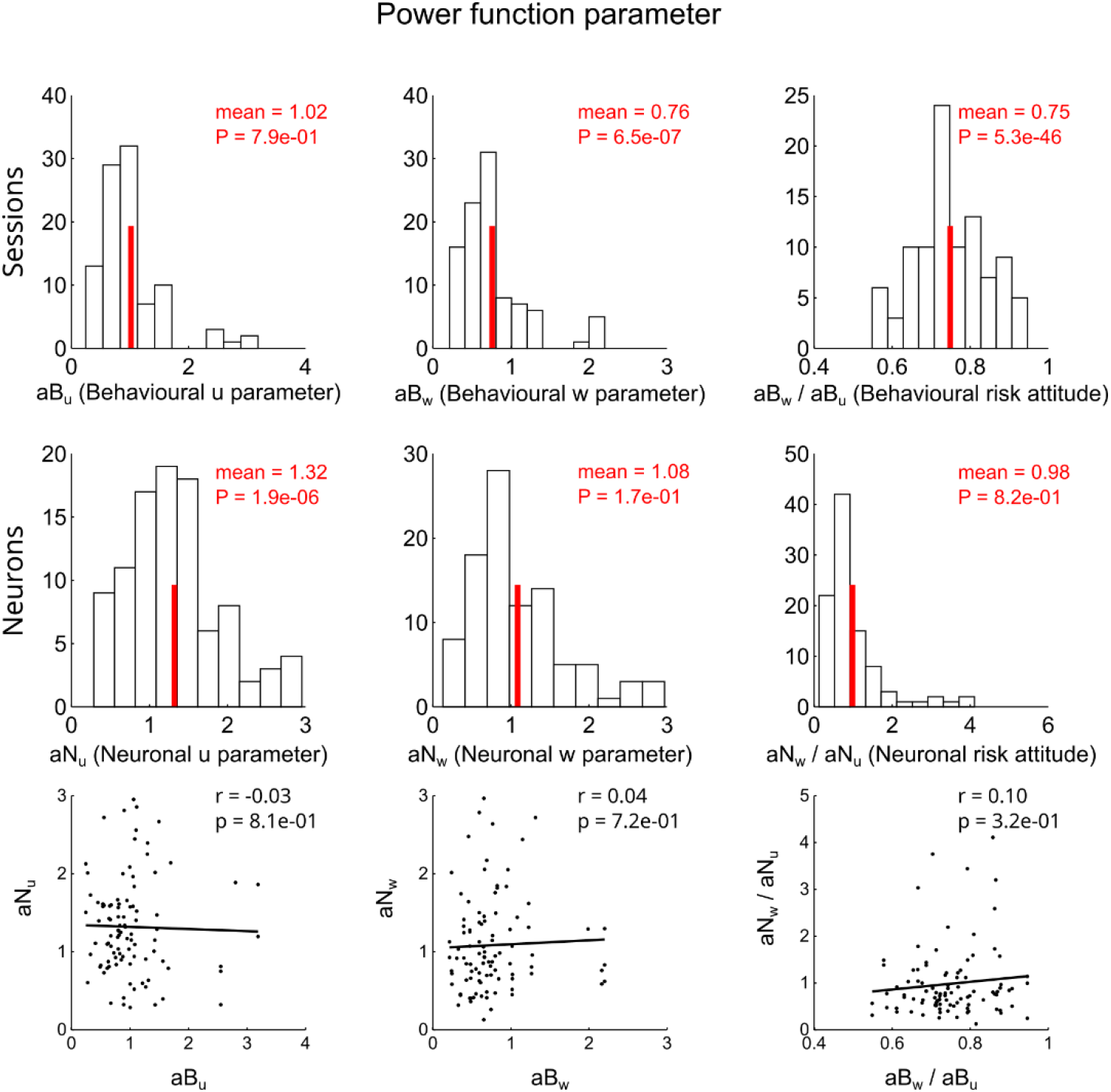
Correlation between neuronal and behavioural risk-attitude measures using power functions. Utility (u parameter) and probability weighting (w parameter) functions were defined as one-parameter power functions (y = x^a^) and fitted to the behavioral data (top) (obtaining best-fitting parameters aB_u_ and aB_w_, respectively) and to the neuronal data recorded in zero-alternative trials (middle) (parameters aN_u_ and aN_w_). The ratio of the two parameters defined a risk-attitude measure that strongly correlated with the alpha measure defined in Raghuraman & Padoa-Schioppa 2014 (Pearson’s correlation: r = 0.86, P = 9.8e-53). This measure was recovered from both behavioral and neuronal data and compared using a correlation analysis (bottom). Data from Monkey B.

**Figure S12.**
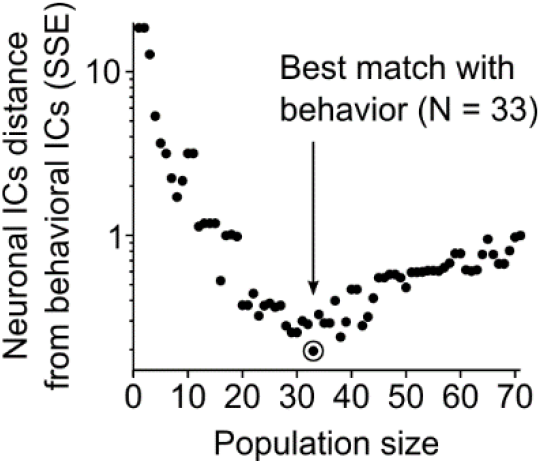
Subpopulations of behavior-incompatible neurons. Reconstruction of the animal’s behavior from combining the two unreliable, behavior-incompatible subpopulations (i.e., those with a significant intercept between neuronal and behavioral preferences, see Figures 4 and 6). The sum of squared errors (SSE) along the probability dimension provided a metric for the difference between neuronal ICs and behavioral ICs. Neurons were sorted based on the absolute value of their intercept, and the SSE was computed for populations of increasing number of neurons (black dots). This distance measure reached a minimum value at a population of N = 33 neurons, thus producing the best fit with behavior. This population was used for the analysis reported in Figure 6G.

**Figure S13.**
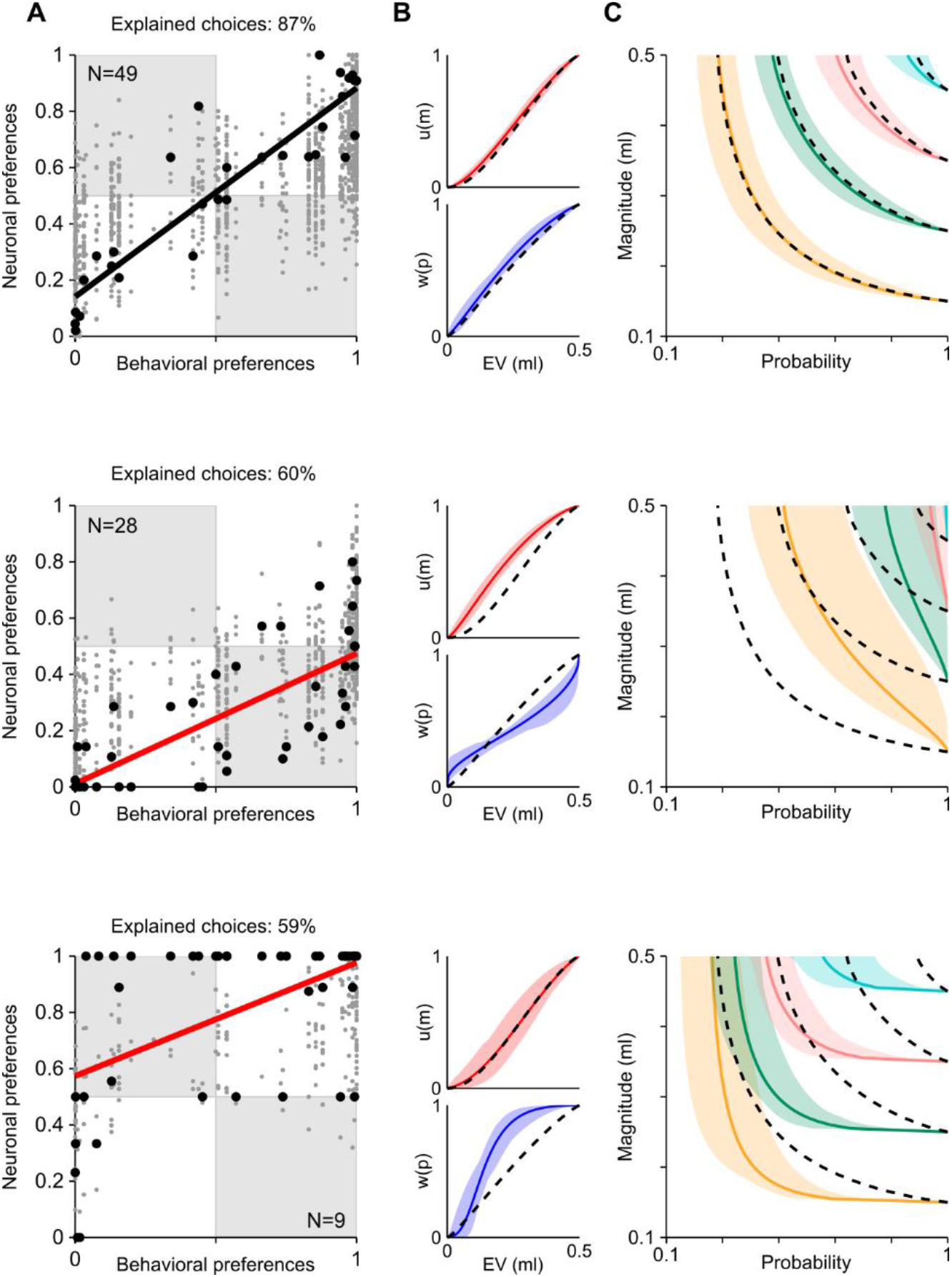
Neuronal population code with stricter neuron selection (P < 0.01). Three subpopulations of positive *m, p* coding neurons with neuronal responses compatible with behavior (top), more risk averse (middle) and more risk seeking (bottom) than behavior. Same analysis as used for Figure 6, using a stricter criterion for the selection of neurons (P < 0.01 instead of P < 0.05; correlation between neuronal and behavioral preferences). The three classes of neurons were reflected in the correlation between neuronal and behavioral preferences (panel A), in the shapes of utility and probability weighting functions (panel B) and in the indifference curves (panel C). Conventions as in Figure 6.

**Figure S14.**
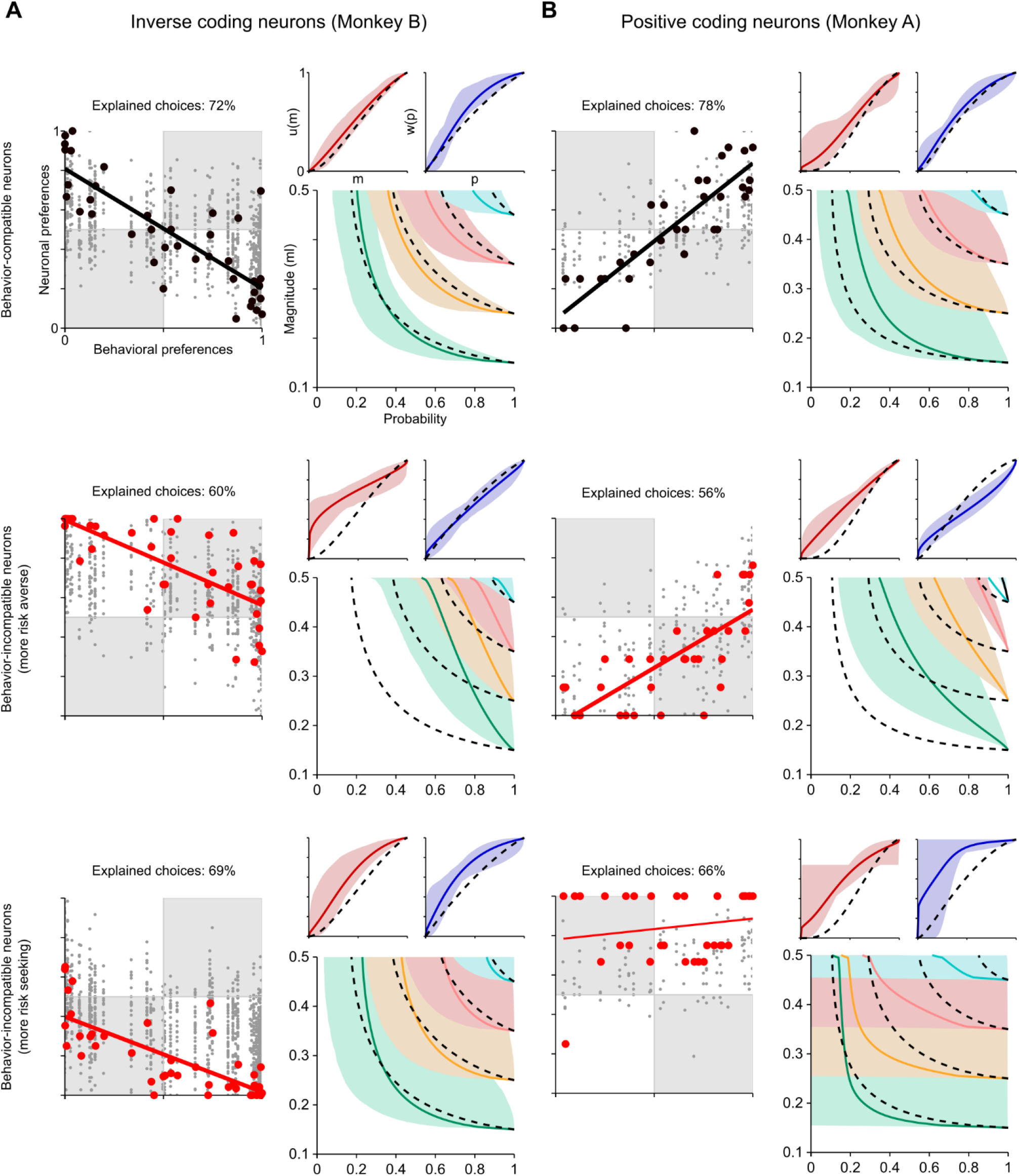
Behavior-compatible and behavior-incompatible subpopulations of positive and inverse coding neurons. (A), (B) Decoded preferences and economic functions for inverse *m, p* coding neurons (panel A) and for positive coding neurons in Monkey A (panel B) confirming and extending the results reported in Figure 6 (conventions and symbols as in Figure 6).

